# Transcutanenous auricular vagus nerve stimulation affects sustained attention in an order-dependent manner

**DOI:** 10.1101/2025.10.07.680948

**Authors:** Christian Wienke, Joshua P. Woller, Agnieszka Zuberer, Alireza Gharabaghi, Tino Zaehle

**Affiliations:** Institute for Medical Psychology, Otto-von-Guericke University, Magdeburg, Germany; Department of Neurology, Section Neuropsychology, Otto-von-Guericke University, Magdeburg, Germany; Research Group “Magdeburger Arbeitsgemeinschaft für Forschung unter Raumfahrt-und Schwerelosigkeitsbedingungen” (MARS); German Centre for Mental Health (DZPG), partner site Halle-Jena-Magdeburg, Germany; Institute for Neuromodulation and Neurotechnology, University Hospital Tübingen, Germany; Max-Planck-Institute for Biological Cybernetics, Tübingen, Germany; Boston University Chobanian and Avedisian School of Medicine, Department of Psychiatry, USA; Centre for Digital Health, University Tübingen, Germany; Centre for Bionic Intelligence Tübingen Stuttgart, University Hospital Tübingen, Germany; German Centre for Mental Health (DZPG), University Hospital Tübingen, Germany; Center for Behavioral Brain Sciences (CBBS), Magdeburg, Germany

## Abstract

**Background:** Sustained attention, the ability to remain focused during a task, relies on Locus coeruleus - Noradrenaline (LC-NA) activity. Noninvasive transcutaneous auricular vagus nerve stimulation (taVNS) is a promising tool to modulate LC-NA activity. We investigated the effects of taVNS on behavioral and electrophysiological markers of sustained attention during a Go-Nogo task.

**Methods:** Forty-four healthy participants (36 female) performed the gradual-onset Continuous Performance Task (gradCPT) on two days ≥ 24 hours apart while EEG was recorded. 500 ms trains of taVNS or Sham stimulation was applied with each stimulus in a pseudorandomized fashion. Brain activity during the task was recorded with EEG. Behavioral and electrophysiological parameters of sustained attention were analyzed using (generalized) linear mixed models.

**Results:** When delivered during the first session, taVNS accelerated reaction times and enhanced fronto-central P1-N1 and parietal N1-P2 amplitudes. Larger P1-N1 amplitudes predicted faster responses, linking early cortical gain to behavioral performance. RT improvements persisted into the next-day sham session, whereas taVNS applied only during the familiar second session was ineffective. However, taVNS also increased RT variability, speaking against a general beneficial effect of taVNS on response automation.

**Conclusion:** We show that taVNS modulates behavioral and electrophysiologial parameters of sustained attention and that this modulation depended on the applied stimulation order. These observations have important implications for future taVNS studies where the order of stimulation is counterbalanced across subjects.

## Introduction

Sustained attention refers to the ability to maintain focus on a task over time (Esterman et al., 2013; Fortenbaugh et al., 2015, 2017). Go-Nogo paradigms, during which participants respond to frequent “Go” stimuli while withholding a response to infrequent “Nogo” stimuli, are commonly employed to assess sustained attention. Failures in response inhibition (commission errors) indicate lapses of attention. Esterman and colleagues (2013) refined this classical paradigm, noting that the abrupt stimulus onsets may exogenously capture attention. They developed the *gradual-onset Continuous Performance Task* (gradCPT) (Esterman et al., 2013) where naturalistic images constantly transition into each other without clear on- and offset. Despite this continuous nature, the task retains a Go-Nogo structure: Frequent city scenes (Go) require a button press while infrequent mountain scenes (Nogo) require response inhibition, allowing the investigation of intra- and interpersonal fluctuations in sustained attention (Esterman et al., 2013).

Neurophysiologically, sustained attention during discrimination tasks (i.e., Go-Nogo tasks) engages right-lateralized prefrontal regions like the right dorsolateral and medial prefrontal cortex (PFC), the right postcentral gyrus and the left posterior ventromedial PFC (Langner & Eickhoff, 2013). Additionally, the dorsal attention network, involving bilateral intraparietal sulcus and frontal eye fields (M. D. Fox et al., 2006), provides top-down control (Vossel et al., 2014). Optimal attentional performance further depends on cortical arousal, as both hypo- and hyperarousal impair attention (Huang & Clewett, 2024).

A key regulator for arousal is the Locus coeruleus - Noradrenaline (LC-NA) system (Aston-Jones & Cohen, 2005; Unsworth & Robison, 2017). The LC, a small brainstem nucleus and main source of cortical noradrenaline (Sara, 2009), projects widely to cortical and subcortical areas, including prefrontal regions implicated in attentional control (Liebe et al., 2022; Szabadi, 2013). It exhibits two distinct activity modes: A tonic background activity is strongly related to arousal (Rajkowski et al., 1994) with low tonic activity during hypoarousal and high tonic activity during hyperarousal (Aston-Jones & Cohen, 2005). Intermediate tonic activity is associated with optimal task performance (Murphy et al., 2011). An additional phasic activity mode is triggered by behaviorally salient or novel stimuli and transiently increases cortical NA levels (Huang & Clewett, 2024). This mode is particularly relevant for attentional engagement, as it selectively amplifies processing in prefrontal and parietal regions critical for attentional control (Devoto et al., 2005). Thus, tonic and phasic LC activity form a dynamic neuromodulatory system that enables flexible adaptation of attention to internal states and external demands (Aston-Jones & Cohen, 2005). The dependency of sustained attention on arousal and LC-NA activity has further been shown in different studies investigating pupil size as proxy for LC-NA activity (Esterman & Rothlein, 2019), where smaller task evoked pupil diameters (i.e., lower LC-NA activity) were associated with longer reaction times (Unsworth & Robison, 2016, 2018).

Pharmacological studies further emphasize NA’s role in sustained attention (Chamberlain & Robbins, 2013). NA-reuptake inhibitors improved sustained attention in animal models (Marshall et al., 2019) and humans (Crockett et al., 2010) (but see Chamberlain et al. (2006) for conflicting results) while a reduction in NA levels impaired sustained attention (Coull et al., 1995; Smith & Nutt, 1996). Notably, LC-NA integrity (measured as contrast ratio in magnetization transfer-weighted MRIs) decreases with age (Liu et al., 2019) and large-scale behavioral data confirm that the ability to sustain attention declines in older adults (Fortenbaugh et al., 2015). Together, these findings underscore the role of the LC–NA system in sustained attention and raise the possibility that external activation of this system may improve attentional performance.

Transcutaneous auricular vagus nerve stimulation (taVNS) is a promising, non-invasive stimulation technique to modulate activity in the LC-NA system. Brief electric pulses are delivered to regions of the auricle innervated by the auricular branch of the vagus nerve (ABVN)(Farmer et al., 2021). The ABVN projects centrally to the nucleus tractus solitarius (NTS), which then projects to several areas including the LC (Butt et al., 2020; Ruffoli et al., 2011). Invasive vagus nerve stimulation in animals increased LC firing and cortical NA release (Hulsey et al., 2017; Manta et al., 2013; Raedt et al., 2011). Correspondingly, human imaging studies have demonstrated increased activation along the vagal afferent path, including the NTS and LC, following taVNS (Borgmann et al., 2021). Optimal stimulation parameters are still under debate but recent studies showed higher effectiveness of short, phasic stimulation trains over tonic or continuous stimulation to engage the LC-NA system (Pervaz et al., 2025). Especially taVNS applied in parallel with stimulus onset improved behavioral accuracy (Wienke et al., 2023), suggesting that stimulation may be more effective when it coincides with ongoing neural activity. The gradCPT specifically has only been used in two stimulation studies so far, applying transcranial magnetic stimulation (TMS) to the frontal eye fields (FEF)(Esterman et al., 2015) and the cerebellar node of the dorsal attention network (DAN)(Esterman et al., 2017). Disrupting activity via TMS in the right FEF selectively increased commission error rate and reaction time variability while being in the zone (Esterman et al., 2015) Facilitating activity in the DAN via cerebellar TMS, on the other hand, reduced commission error rate but had no effect on reaction time parameters (Esterman et al., 2017)

On an electrophysiological level, the amplitude of the fronto-central N2 component of the event-related potential (ERP) is larger for Nogo trials and has been associated with cognitive control and the inhibition of a planned response (Folstein & Van Petten, 2008). Additionally, the centro-parietal P3 component is associated with the allocation of attentional resources (Polich, 2007) and has been linked to phasic LC-NA activity (Nieuwenhuis et al., 2005).

These findings underscore the potential of taVNS to enhance sustained attention during the gradCPT. However, several important questions remain. The gradCPT has been extensively studied in behavioral (Fortenbaugh et al., 2015; Yamashita et al., 2021) and neuroimaging studies (Esterman et al., 2013; Fortenbaugh et al., 2018; Kucyi, Esterman, et al., 2016; Zuberer et al., 2021). Electrophysiological investigations, on the other hand, are scarce and have predominantly relied on intracranial recordings in clinical populations (Akkol et al., 2021; Kucyi, Esterman, et al., 2016). To date, no study has systematically examined EEG correlates of the gradCPT in healthy adults. TaVNS applications in Go-Nogo paradigms are also limited. Camargo et al. (2024), for example, reported taVNS specific increases in N2 amplitudes during Nogo trials after controlling for handedness, mood and fatigue levels but found no modulation of the P3. In contrast, taVNS related reductions of the N2 amplitude in an Executive Reaction Time Test (Pihlaja et al., 2020) and a Simon conflict task (Fischer et al., 2018), both requiring cognitive control and response inhibition, have been reported. For the P3, increased amplitudes following taVNS in oddball paradigms (Chen et al., 2023; Giraudier et al., 2024; Gurtubay et al., 2023) are reported although null findings also exist (D’Agostini et al., 2023; Fischer et al., 2018). Behavioral results from Go-Nogo tasks during taVNS are likewise rare. While some studies report improved accuracy following taVNS (Keute et al., 2020; Zhu et al., 2024), others found no effect (Camargo et al., 2024) or even impaired performance (Bömmer et al., 2024). Studies employing other paradigms demonstrated faster reaction times during an auditory oddball (Gurtubay et al., 2023) and a visual dot probe task (Chen et al., 2023) as well as improved accuracy in an emotional Stroop task (Wienke et al., 2023).

Here we investigate whether canonical ERP components associated with response inhibition, namely the N2 (Folstein & Van Petten, 2008) and the P3 (Polich, 2007), can be reliably elicited during the gradCPT despite its continuous stimulus transition. Furthermore, we examined whether taVNS modulates these electrophysiological markers as well as behavioral performance in a context-dependent manner. Based on the involvement of phasic LC–NA activity in sustained attention and the effects of taVNS on noradrenergic signaling, we hypothesized that taVNS would improve behavioral performance during the gradCPT. Behaviorally, we expected faster and more stable reaction times; electrophysiologically, we anticipated robust, early ERP components resembling the canonical N2 and P3 responses that would be modulated by trial type (Nogo > Go) and enhanced under taVNS relative to sham stimulation

## Materials and Methods

### Study sample

A total of 44 participants (36 female) aged between 18 and 30 (*M* = 21.7, *SD* = 3.2) took part in this study. Sample size was determined to match previous taVNS studies where behavioral and electrophysiological data were recorded (D’Agostini et al., 2023; Wienke et al., 2023). All participants were right handed, had normal or corrected to normal vision and reported no history of neurological or psychological disorders. Exclusion criteria were the following: electrically active implants (e.g. pacemaker, cochlear implants), head injuries within the past 6 months, history of psychological or neurological disorders, psychoactive medication, arrhythmia, pregnancy. Written informed consent was obtained prior to participation. Recordings took place at the Otto-von-Guericke University, Magdeburg and the study was approved by the local ethics committee (‘Ethical committee of the Otto-von-Guericke University Magdeburg’; Az: 95/18). Participation was reimbursed with course credit or money.

### Experimental procedure and task

We employed a single-blind, within-subject, counterbalanced design consisting of two sessions. In each session, participants performed the gradual-onset continuous performance task (gradCPT; adapted from Kucyi et al. (2016)) while receiving either active taVNS or sham stimulation. The order of stimulation (taVNS first and sham first) was pseudo-randomized and balanced across participants. Sessions were spaced at least 24h and at most 7 days apart (*M* = 2.98, *SD* = 2.03, range: 1-7 days) with no significant difference between both order groups (*t_42_* = -0.81, *p* = 0.42).

Following EEG and stimulation setup, participants performed four blocks of the gradCPT per session (Fig. 1A). Stimuli were presented using Matlab R2018b (The MathWorks, Inc., Natick, Massachusetts, United States) and the Psychtoolbox 3 (Brainard, 1997; Kleiner et al., 2007; Pelli, 1997). Each block lasted approximately 9 minutes (Fig. 1B) and consisted of 375 trials. Stimuli were round, greyscale images of city scenes (Go trials, 90%) and mountain scenes (Nogo trials, 10%) adapted from the SUN database (Xiao et al., 2010). Images gradually faded into each other over a 1.3 s transition (Fig. 1C). Participants were instructed to press the space bar in response to city scenes and to withhold a response for mountain scenes. TaVNS or sham stimulation (see below) was applied at the onset of each new stimulus, i.e., when a new scene started to fade in (Fig. 1C). Each session started with two training runs of 1 min duration to familiarize participants with the task. In the first training run, transition time between stimuli was double (i.e. 2.6 s) while the second training run used the same transition duration as the actual experimental blocks. No stimulation was applied during these training runs. Participants were encouraged to respond as accurately and fast as possible.

**Fig. 1.**
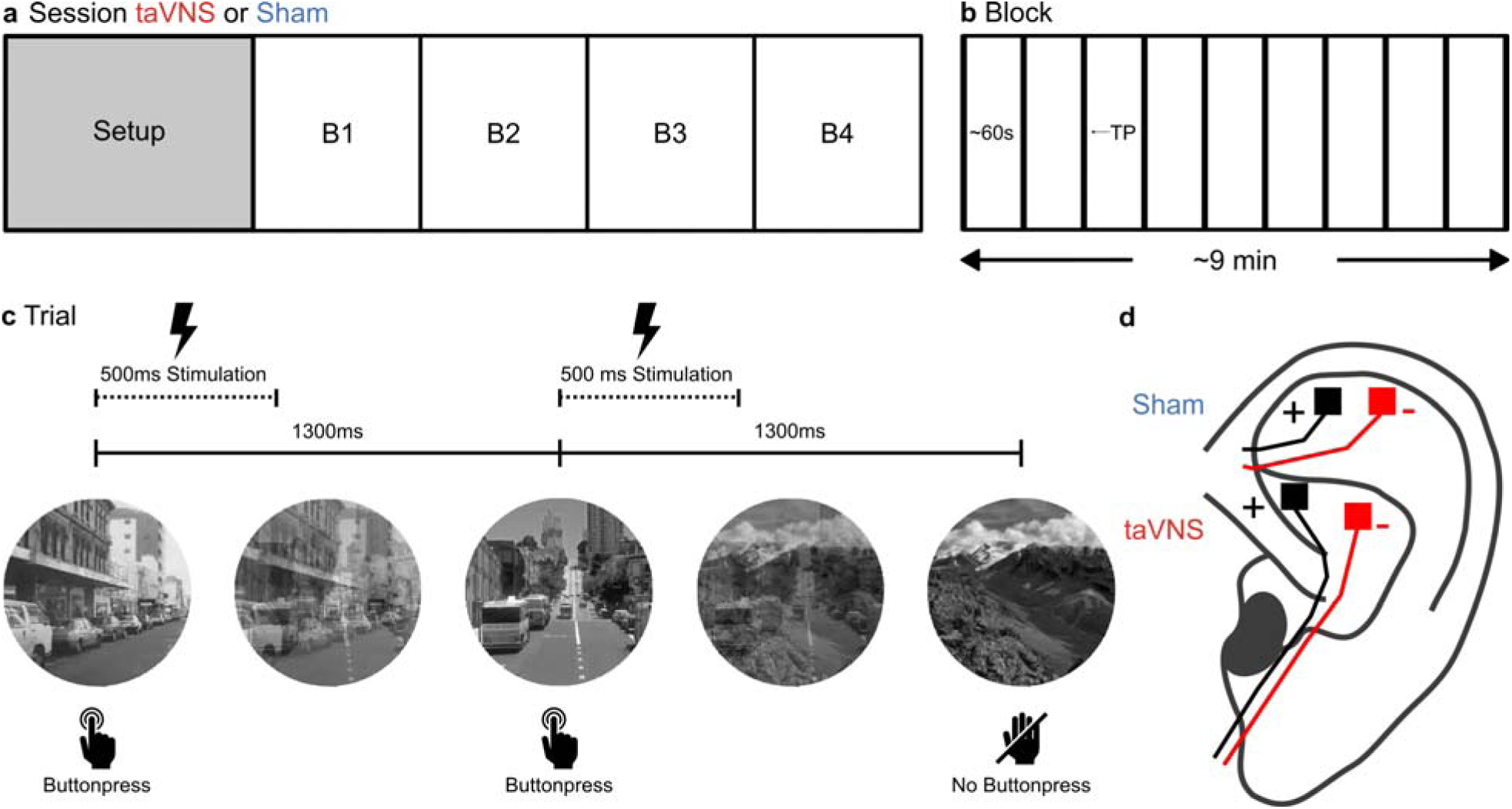
Experimental design and stimulation protocol. A) Session structure: Each participant completed two sessions in pseudorandomized order, receiving either transcutaneous auricular vagus nerve stimulation (taVNS) or sham stimulation. Following EEG setup, each session comprised four consecutive blocks of the gradual-onset continuous performance task (gradCPT). B) Block structure: Each block lasted ∼9 minutes and contained 375 trials and 9 intermittent thought probes (TP) assessing subjective task focus. C) Task trial timeline: Stimuli consisted of round grayscale city (frequent Go) and mountain (rare Nogo) scenes that gradually transitioned every 1300 ms. Participants pressed a button for city scenes and withheld a response for mountain scenes. A 500 ms train of electrical stimulation was administered time-locked to each stimulus onset in both, the taVNS and sham condition. D) Stimulation montage: For taVNS, electrodes were placed on the cymba conchae of the left ear. For sham, electrodes were placed on the scapha of the same ear, avoiding vagal innervation.

Each block contained 9 thought probes presented approximately once per minute (Fig. 1B). During these thought probes, participants were first asked to rate the extent to which their attention was focused on the task or elsewhere. A visual rating scale (VRS) was displayed, anchored with “only task” and “only something else”. Participants moved a cursor along the scale using the arrow keys and confirmed their selection with the space bar. A second question then assessed meta-awareness (i.e., to what extent they were aware about their focus of attention) using a similar VRS anchored with “completely aware” and “completely unaware”. Responses on both VRS were recorded on a 0-100 scale, though values were not displayed to participants. The task resumed immediately after the second response. Participants were allowed self-paced breaks between blocks.

At the end of each session, participants completed two questionnaires. The first assessed potential adverse effects of stimulation (e.g. headache, nausea, dizziness, skin irritation) on a 4-point scale from 0 (none) to 3 (strong). The second questionnaire asked participants to rate their confidence in monitoring their attentional focus during the thought probes, followed by ratings of how often periods of reduced task focus were due to external distractions, internal task-related thoughts (e.g., about performance or strategy), or task-unrelated thoughts (e.g., mind-wandering). These items were rated on a 7-point scale from 1 (completely not confident / never) to 7 (completely confident / always).

### Electrical stimulation

Stimulation was applied as trains of monophasic, 200 µs long square wave pulses at 30 Hz for 500 ms coinciding with the onset of each picture as in recent studies (Rufener et al., 2023; Wienke et al., 2023). For active taVNS, electrodes were attached to the cymba conchae of the left auricle. For the sham stimulation, electrodes were attached to the scapha of the left auricle (Fig. 1D). This area was chosen because the high heterogeneity of results in sham controlled taVNS studies questioned the validity of the ear lobe as a suitable control condition (Borges et al., 2021; Rangon, 2018). Cakmak (2019) argued that a lower number of sympathetic nerve fibers in the ear lobe needs to be considered. Upper parts of the auricle, on the other hand, exhibit similar densities of sympathetic fibers as the cymba conchae (Cakmak, 2019). In both cases, Ag/AgCl surface electrodes (Ambu Neuroline 700, Ambu, Germany), cut to a size of 4×4 mm and placed ca. 1 cm apart, were used. Prior to electrode placement, the skin was cleaned using disinfectant alcohol. A small amount of conductive paste (lic2 low impedance cream, www.cnsac-medshop.com) was used to assure proper conductance. Pulses were delivered by a Digitimer DS7A (Welwyn Garden City, UK) connected to the two electrodes. The Digitimer was controlled via custom Matlab scripts using the Data Acquisition Toolbox and a digital/analog converter (DAC; NI USB-6212, National Instruments, Austin, TX, USA). Stimulation intensity was set to 2 mA for all participants and tested before starting the experiment by applying single stimulation trains of 500 ms duration. If 2 mA felt uncomfortable, the intensity was reduced in 0.1 mA steps until no longer uncomfortable. The average stimulation intensity was 1.96 mA for sham (*SD:* 0.15) and 1.91 mA for taVNS (*SD* = 0.27). There was no statistical difference between conditions (*t_43_*: 1.2, *p* = 0.24).

### EEG recording and processing

The EEG was continuously recorded throughout the task from 34 Ag/AgCl electrodes, placed according to the international 10-10 system, via a BrainAmp amplifier (www.brainproducts.com) with 1000 Hz sampling frequency. The online reference was placed on the nose, the ground electrode at AFz. Impedance for all electrodes was kept below 10 k . A salt free, abrasive, conductive paste (Abralyt 2000, Brain Products, Gilching, Germany) was used to ensure conductivity. A vertical electrooculogram (EOG) was recorded from below the right eye. Remaining electrodes were Fpz, Fp1, Fp2, F1, F2, F3, F4, F7, F8, FT9, FT10, FCz, Cz, C3, C4, CP5, CP6, T7, T8, TP9, TP10, Pz, P7, P8, POz, PO3, PO4, PO8, PO9, O1, O2, Iz, left and right mastoid.

Offline, the stimulation artifacts were removed from the EEG by applying a spatial filter derived via generalized eigendecomposition (GED) (Cohen, 2022; Haslacher et al., 2021; Woller et al., 2024), implemented in Python using MNE, SciPy, and NumPy. GED, similar to ICA, extracts multivariate components from the data, but here we contrasted covariances around stimulation pulses with baseline periods, allowing for an a priori tuning of the decomposition towards artifact components. In the present data, 1–3 components were sufficient to attenuate the artifact. The remaining preprocessing of the EEG data was performed in Matlab R2024b using the Fieldtrip toolbox (Oostenveld et al., 2011). Continuous data after removal of the stimulation artefact were filtered between 1 and 30 Hz using a bidirectional infinite impulse response Butterworth filter. An additional Band-Stop filter between 49 and 51 Hz was applied to suppress line noise. Data were then downsampled to 200 Hz and remaining artefacts, e.g. from blinks, were removed using an ICA approach (Infomax algorithm). Afterwards the continuous EEG signal was segmented into epochs from -500 to 1200 ms relative to stimulus onset. Epochs exceeding absolute amplitudes of 200 µV peak-to-peak in a 200 ms window were automatically excluded. One subject was excluded from the analysis due to more than 50% of removed trials. In the remaining participants, an average number of 5.69 Go-trials (*SD:* 8.23, range: 0-38) and an average number of 0.65 Nogo-trials (*SD:* 1.17, range: 0-7) was excluded. The amount of excluded trials between sessions did neither differ for go-trials (*t_42_* = -0.75, *p* = 0.46) nor for nogo-trials (*t_42_* = 0.97, *p* = 0.34).

Event related potentials (ERPs) were computed from correct Go trials for each block, stimulation condition and session. For the correct Nogo condition, blocks 1 and 2 as well as 3 and 4 were combined, to obtain a sufficient number of trials due to the low probability of Nogo trials. For the incorrect Nogo condition, all 4 blocks were combined for the same reason. No baseline correction was performed, since the gradCPT does not provide a suitable, stimulus free period immediately prior to each stimulus onset. Peak-to-peak amplitudes were instead extracted as they are independent of a baseline correction (McClaskey et al., 2018). In the Go condition, the Grand Average ERP across all blocks, sessions and stimulation conditions showed an early positive as well as a later negative deflection primarily at fronto-central electrodes FCz and Cz (Fig. 2A). The first positive peak (“P1”) was defined as the most positive local maximum between 200 and 600 ms after stimulus onset. The subsequent negative peak (“N1”) was defined as the most negative local minimum occurring within 400 ms after the P1 peak. The peak-to-peak amplitude was calculated as the difference between these two peaks. In addition, a second prominent positive deflection was observed, particularly at electrode Pz (Fig. 2C). The peak-to-peak amplitude was calculated as difference between the most negative peak occurring between 300 and 700 ms post-stimulus and the most positive peak occurring within 500 ms after this negative peak. In the grand average ERP of the Nogo condition (Fig. 3A), we observed pronounced negative and positive deflections with a more parietal distribution centered around electrode Pz. As with the Go condition, peak-to-peak amplitudes between the negative and positive deflections at electrode Pz were computed for each stimulation condition, session, and the combined block pairs (blocks 1+2 and blocks 3+4). For the incorrect Nogo trials the N1-P2 amplitude was determined in the same way for both sessions and stimulation conditions.

**Fig. 2.**
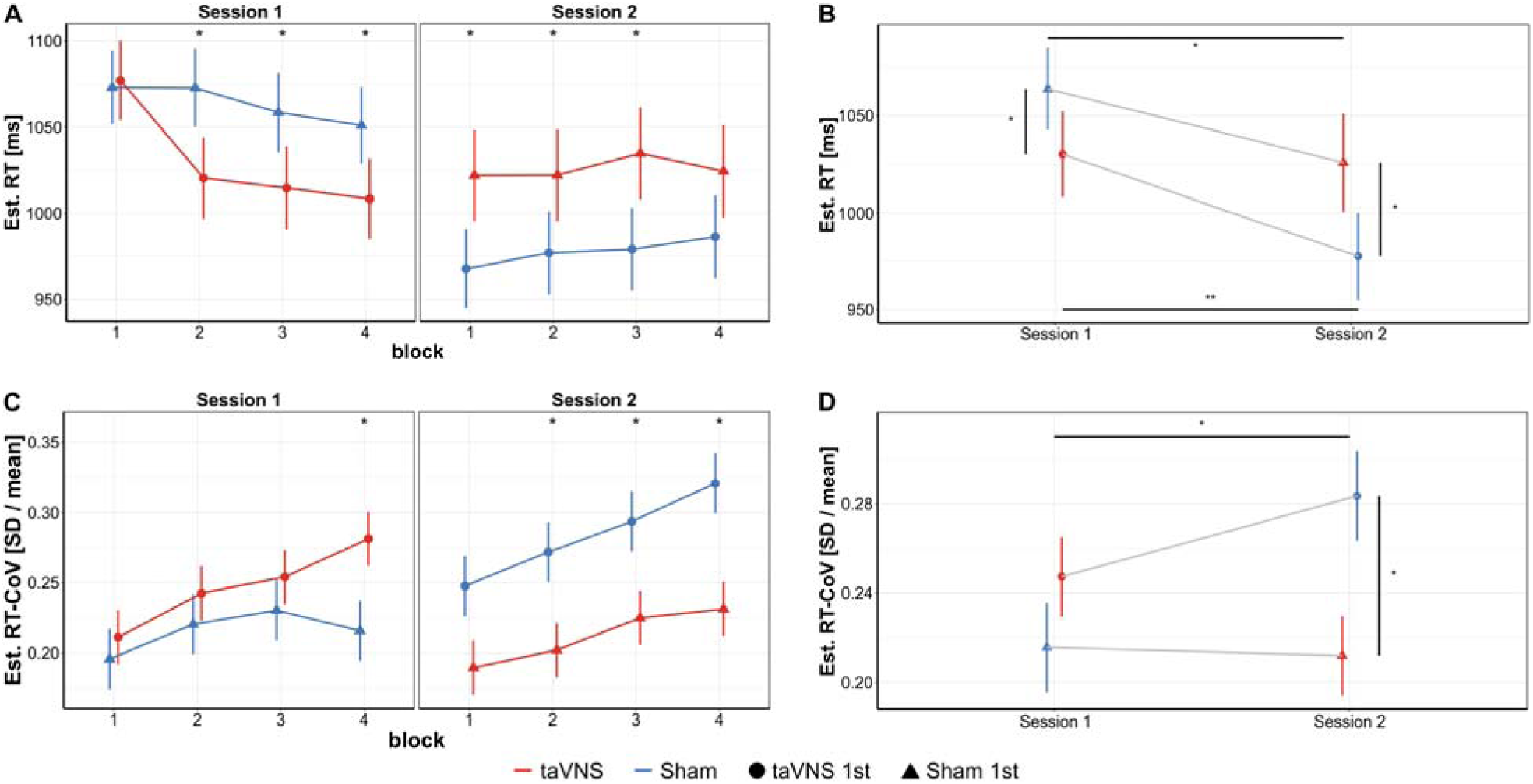
TaVNS improves reaction time (RT) only when the task is novel but increases variability. A) During the first exposure to the gradCPT (Session 1, left panel) taVNS (red) produced a rapid RT drop after Block 1, whereas sham (blue) showed no change. Importantly, both groups started with comparable RTs in Block 1. When the same participants returned ≥24 h later (Session 2, right panel) the taVNS-first group (circles) retained faster RTs during sham stimulation (Blocks 1 to 3), indicating a persistent benefit, while the sham-first group (triangles) still showed no improvement. B) RT gains within Sessions and from Session 1 to Session 2 were larger for the taVNS-first group than for the sham-first group. C) RT-CoV in Session 1 continouosly increased over the course of the session for the taVNS first group whereas the sham-first group reached a plateau between block 3 and 4. In Session 2, the taVNS-first group again retained higher RT-CoV values. D) Averaged across blocks, the taVNS first group showed a significant increase in RT-CoV from Session 1 to Session 2, while the sham-first group showed no increase. Error bars denote s.e.m.; *p < 0.05, **p < 0.01 (FDR-corrected)

**Fig. 3.**
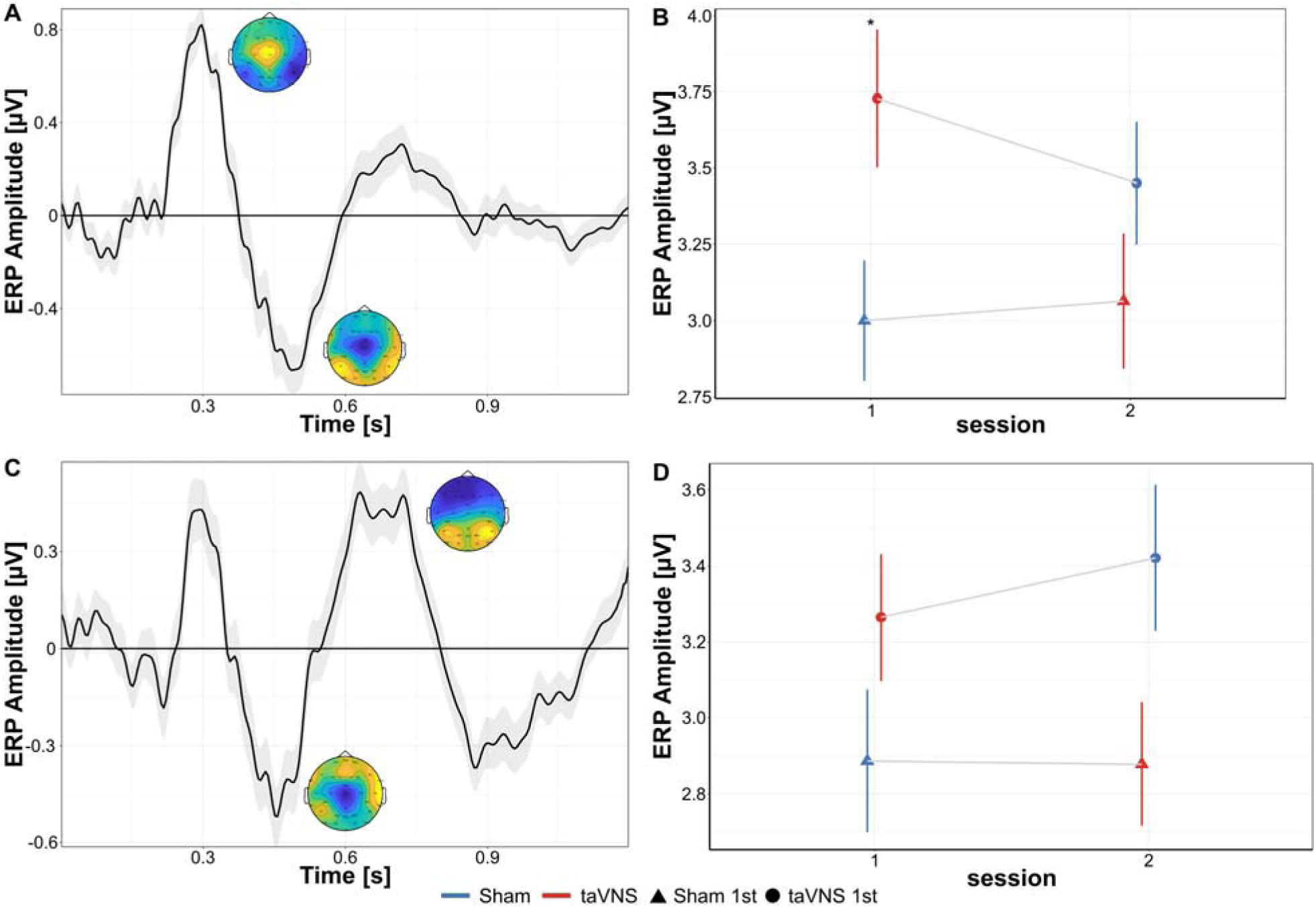
TaVNS amplifies early sensory–attentional ERPs during the novel session only. A) Fronto-central waveform. Grand-average event-related potential (ERP) for correct Go trials at FCz + Cz shows an initial positive peak (∼300 ms) followed by a negative deflection (∼500 ms); insets depict scalp voltage at each peak. B) P1–N1 amplitude (FCz + Cz). A significant stimulation - session interaction revealed larger peak-to-peak amplitudes when taVNS was delivered in Session 1 (red) relative to sham (blue); no difference emerged in Session 2. C) Parietal waveform. Grand-average ERP at Pz exhibits a pronounced second positive peak. D) N1–P2 amplitude (Pz). The same stimulation - session interaction was observed, although post-hoc contrasts did not reach FDR-corrected significance. Shaded areas (A, C) and error bars (B, D) denote s.e.m.; P < 0.05, **P < 0.001 (FDR-corrected).

### Behavioral parameters

*Reaction times (RTs)* were calculated as in previous studies (Esterman et al., 2013; Kucyi, Esterman, et al., 2016) relative to onset of each image transition and used to define trials into *in the zone* and *out of the zone* based on the variance time course (Esterman et al., 2013). A detailed description can be found in the supplementary material. The *Reaction time coefficient of variation (RT-CoV)* was calculated as a marker of response stability (MacDonald et al., 2006) by dividing the standard deviation of correct Go trials by their mean (Fortenbaugh et al., 2015) for each block and session within each participant. As an index of *discriminatory ability*, *d’* was computed based on hit rate and false alarm rate (Fortenbaugh et al., 2015, 2018) for each block and session. As a complementary marker of discriminatory ability, the parameter *A* was calculated as it does not depend on the artificial insertion of errors and an underlying normal distribution (Zhang & Mueller, 2005). A detailed description, including the formulas used for calculation, can be found in the supplementary material. We also determined the number of *in the zone* trials based on the variance time course of both concatenated sessions. This allowed us to analyze whether taVNS affected task engagement within participants.

### Statistical Analyses

Statistical analyses were performed using R 4.3.3 (R Core Team, 2024) and RStudio 2021.09.1 (RStudio Team, 2021). Linear mixed effect models (LMMs) and generalized linear mixed effect models (GLMMs) were fitted with the *lme4*-package (Bates et al., 2015). For each model, we initially specified a maximal random effects structure justified by the design (Barr et al., 2013). If the maximal model did not converge, the random-effects structure was reduced step wise. Competing nested models were compared using likelihood ratio tests (a = 0.05) and the AIC. The best-fitting converging model was retained for inference. Model comparisons were performed using the *anova()* function. Final model formulas are provided in the supplementary material for brevity. Significance of predictors in the final models were estimated using the *Anova()* function from the *car*-package (J. Fox & Weisberg, 2019) which calculates F-tests for fixed effects and returns X^2^ statistics. Post-hoc tests were conducted with the *emmeans* package (Lenth, 2022), using pairwise comparisons and applying a False Discovery Rate (FDR) correction based on the Benjamini–Hochberg procedure to account for multiple comparisons. Model assumptions were assessed using graphical residual diagnostics. Histograms and QQ-Plots were used to assess normality and residuals vs. fitted values as well as standardized residuals were used to assess homoscedasticity and potential outliers.

End-of-session questionnaire responses were analyzed using Wilcoxon signed-rank tests to compare taVNS and sham conditions. For all other dependent measures, the main predictor of interest was *stimulation* (taVNS vs. sham). *Block* (1–4) and *session* (1 vs. 2) were included to account for potential effects of e.g. tiredness or training within and across sessions. Interactions included all two- and the three-way interaction. For behavioral data, the categorical covariate *zone* (in the zone vs. out of the zone) was included to account for different task engagement. RTs were analyzed using a GLMM with gamma distribution and identity link function (Lo & Andrews, 2015) while for the RT-CoV a GLMM with gamma distribution and log-link function was used. To make sure that potential effects were not just scaling artifacts due to differences in RT, we included the person-centered mean RT as covariate in this model to control for changes in RT (Wagenmakers & Brown, 2007). The number of *in the zone* trials across sessions was analyzed using a negative binomial GLMM due to overdispersion. Subjective task focus as measured by the thought probes, discriminatory ability parameters (*d’*, *A*), and ERP peak-to-peak amplitudes were analyzed using LMMs. Random effects in all models included random intercepts across participants. Final model formulas are provided in the supplementary material.

## Results

### Adverse stimulation effects

Participants did not report any serious adverse effects in response to either taVNS or sham stimulation. Ratings of minor side effects (e.g., tingling, headache, dizziness) were low and did not differ significantly between stimulation conditions (all *p*’s > 0.24). Additionally, mind wandering was reported more often as cause for distraction, although no differences between taVNS and sham were observed. Details can be found in tables S1 and S2.

### General characteristics of the gradCPT

We first report general effects of the gradCPT on behavioral and ERP parameters. Behavioral analyses on reaction time (RT) and discriminatory ability (*d’* and *A*) aimed to replicate and extend the original work (Esterman et al., 2013). We also analyzed the *Reaction Time Coefficient of Variation* (RT-CoV) as additional marker of sustained attention (Yamashita et al., 2021). Predictors include the *block* and *session* number as well as the *zone* of attention, estimated from the response time variability (Esterman et al., 2013). ERP parameters focus on early, fronto-central and later parietal components defined from the Grand Average ERP (see Methods section). In accordance with the original work (Esterman et al., 2013), *RTs* were significantly modified by the *zone* of attention (X*^2^_(1)_* = 150.39, *p* < 0.001). Participants responded faster during *in the zone* trials (*M* = 992 ms, *SE* = 21 ms) compared to *out of the zone* trials (*M* = 1057 ms, *SE* = 20 ms). An additional main effect of *session* was observed (X*^2^_(1)_* = 51.706, *p* < 0.001) with shorter RTs in the second (*M:* 1002 ms, *SE:* 22 ms) than during the first session (*M:* 1047 ms, *SE:* 20 ms). A *block* - *session* interaction (X*^2^_(3)_* = 8.814, *p* = 0.032) indicated longest RTs in block 1 of session 1 which declined in block 2 (*p_corrected_* = 0.01), 3 (*p_corrected_*< 0.001), and 4 (*p_corrected_* < 0.001) but no differences between blocks in the second session (all *p*’s > 0.55).

For the RT-CoV, the predictor *zone* was not included in the model to avoid circularity, as it was determined from a different parameter of reaction time variability. Here, we observed a significant main effect of *session* (X*^2^_(1)_* = 4.6, *p* = 0.03), *block* (X*^2^_(3)_* = 10.51, *p* = 0.01) and a significant *block* - *session* interaction (X*^2^_(3)_* = 10.12, *p* = 0.02). While RT-CoV generally increased from block 1 (*M:* 0.21, *SE:* 0.01) to block 4 (*M:* 0.26, *SE:* 0.01, *p_corrected_*< 0.001), and from session 1 (*M:* 0.23, *SE:* 0.013) to session 2 (*M:* 0.25, *SE:* 0.013), this increase reached a plateau in session 1 with no significant difference between block 2 and 3 (*p_corrected_*= 0.13) or between block 3 and 4 (*p_corrected_* = 0.36). In Session 2, however, RT-CoV increased continuously with significant differences between block 2 and 3 (*p_corrected_* = 0.02) as well as between block 3 and 4 (*p_corrected_* = 0.05).

The discriminatory parameters *d’* and *A* were only modulated by the *zone* of attention (*d’*: X*^2^_(1)_* = 54.12, *p* < 0.001; *A*: X*^2^_(1)_* = 24.78, *p* < 0.001). While being *in the zone*, *d’* was higher (*M:* 4.48, *SE:* 0.07) compared to *out of the zone* trials (*M:* 4.2, *SE:* 0.07). Similarly, *A* was higher *in the zone* (*M:* 0.993, *SE:* 0.002) than *out of the zone* (*M:* 0.986, *SE:* 0.002).

Subjective task focus, measured by the interspersed thought probes, was modulated by the predictor *block* (X*^2^_(3)_* = 70.185, *p* < 0.001) and declined significantly from block 1 (*M:* 76.15, *SE:* 2.49) to block 4 (*M:* 61.2, *SE:* 2.91, *p* < 0.001, Fig. S1).

Finally, self-reported meta awareness was similarly affected by the block within a session (X*^2^_(3)_* = 129.94, *p* < 0.001) but also by a *block* - *session* interaction (X*^2^_(3)_* = 18.92, *p* < 0.001). Meta-awareness again declined from block 1 (*M:* 75.16, *SE:* 62.1, *SE:* 2.96, *p_corrected_* < 0.001) but this decline was stronger and more continuous in session 1 (Fig. S1)

For the ERP components during correct Go trials, the parietal (Pz) N1-P2 amplitude was modulated by the predictor *session* (X*^2^_(1)_* = 7.69, *p* = 0.01) with higher amplitudes in the second (*M:* 3.15, *SE:* 0.13) compared to the first session (*M:* 3.08, *SE:* 0.12). For the correct Nogo trials, ERP amplitudes were considerably larger than during Go trials (Fig. 4A). Apart from that, N1-P2 amplitudes at Pz were not modulated by a general *session* or *block* effect (all *p*’s > 0.11). N1-P2 amplitudes from incorrect Nogo trials were significantly lower compared to correct Nogo trials (X*^2^_(1)_* = 109.76, *p* < 0.001) and followed a more central topography similar to the Go trials (Fig. 3A for comparison). Interestingly, we also observed an early positive deflection peaking at around 340 ms after stimulus onset at central electrodes, resembling the P1 peak during Go trials (Fig. 4C). No effect of *stimulation*, *session*, or an interaction between these two (all *p*’s > 0.38) was observed. However, as the number of trials available for this ERP was small, these results need to be interpreted with care.

**Fig. 4.**
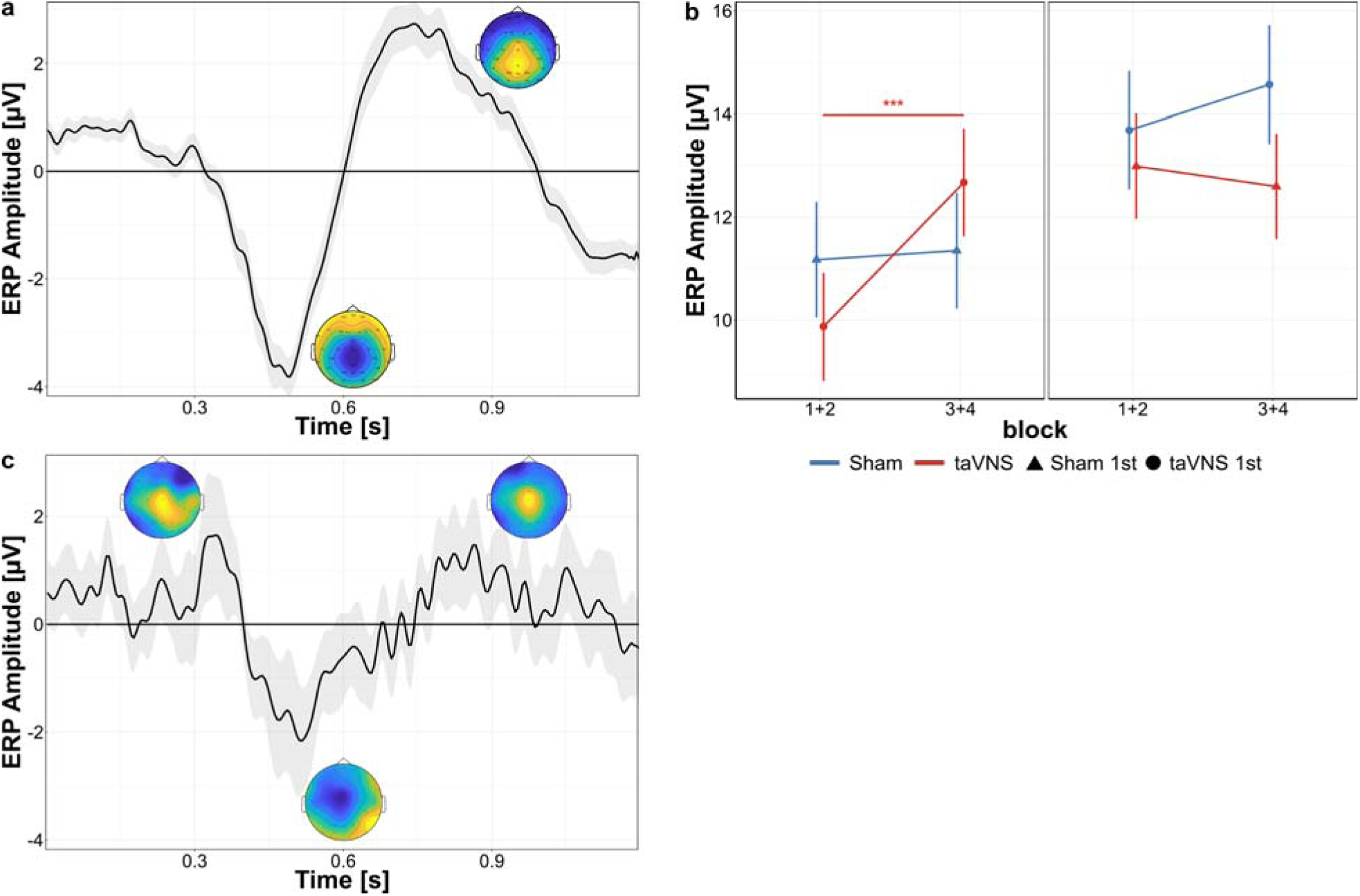
TaVNS modulates parietal Nogo ERPs during the novel session. A) Parietal waveform. Grand-average event-related potential (ERP) for correct Nogo trials (electrode Pz) shows a large negative peak (∼650 ms) followed by a broad positive deflection (∼800–900 ms); insets depict scalp voltage at these extrema. B) Block-wise N1–P2 amplitude (Pz). A three-way stimulation - session - block interaction revealed that participants who received taVNS in Session 1 (red) exhibited a pronounced increase in N1–P2 amplitude from the first half of the session (blocks 1 + 2) to the second half (blocks 3 + 4). No within-session change was observed in the sham-first group (blue), and direct taVNS-vs-sham contrasts were not significant in either half. C) Fronto-central waveform. For completeness, the grand-average Nogo ERP at FCz + Cz is displayed; no stimulation effect emerged here. Shaded areas (A, C) and error bars (B) denote s.e.m.; ***P < 0.001 (FDR-corrected).

### Stimulation specific effects on gradCPT characteristics

Next, we focus on stimulation specific effects during the gradCPT. Visual inspection of the RT data (Fig. 2A) suggested that taVNS only had an effect when applied in the first session. The corresponding GLMM indicated that RTs were indeed modulated by a *stimulation*-*block*-*session* interaction ( *^2^_(3)_* = 14.497, *p* = 0.002, Fig. 2A and table S3). Post-hoc comparisons between sham and taVNS per block and session revealed significant differences during block 2 (*p_corrected_* = 0.03), 3 (*p_corrected_* = 0.04), and 4 (*p_corrected_* = 0.04) of the first session as well as during block 1 (*p_corrected_* = 0.03), 2 (*p_corrected_* = 0.04), and 3 (*p_corrected_*= 0.03) of the second session. Importantly, no differences in RT were observed during the first block of the first session when all participants were naive to the task (*p_corrected_* = 0.78). These results suggest that the effect of taVNS depended on whether it was applied during the first, novel session.This was further demonstrated by the significant *stimulation*-*session* interaction ( *^2^_(3)_* = 4.707, *p* = 0.03, Figure 2B). RTs differed between taVNS and sham in both, session 1 (*p_corrected_* = 0.03) and session 2 (*p_corrected_* = 0.01). The more interesting aspect of this interaction, however, is the effect within the taVNS-first and the sham-first group. Participants receiving taVNS in the first session had lower RTs (*M* = 1030, *SE* = 22) compared to participants receiving sham stimulation (*M* = 1064, *SE* = 21). These participants also showed shorter RTs in the second session (*M* = 978, *SE* = 23), where they then received sham stimulation, compared to participants receiving taVNS in the second session (*M* = 1026, *SE* = 25). The RT improvement from the first to the second session was stronger for the taVNS-first group (−53 ms, *p_corrected_* = 0.003) than for the sham-first group (−38 ms, *p_corrected_* = 0.02, Figure 2B). Importantly, the significant *stimulation* - *session* interaction reflects a reversal of the direction of the taVNS - sham contrast across sessions, which in turn results from the different orders of stimulation. Since taVNS and sham were each applied in both sessions but to different subject groups, this statistical interaction should not be interpreted as a within-subject interaction effect. Instead, it reflects a main effect of stimulation order that manifests as a crossover in the taVNS–sham contrast between sessions.

For the RT-CoV, we observed a similar *stimulation*-*block*-*session* interaction (X*^2^_(3)_* = 13.193, *p* = 0.004, Fig. 2 and table S4). In session 1, RT-CoV increased over blocks with a significant difference between taVNS and sham in block 4 (*p_corrected_*= 0.05). In session 2, the taVNS first group exhibited overall higher variability with significant differences between taVNS and sham in block 2 (*p_corrected_*= 0.04), 3 (*p_corrected_* = 0.04), and 4 (*p_corrected_*= 0.02). Although no specific *stimulation*-*session* interaction was observed in the model (X*^2^_(1)_* = 1.772, *p* = 0.183), exploratory post-hoc comparisons showed a significant increase in RT-CoV from session 1 to session 2 only in the taVNS first group (*p_corrected_*= 0.003) but not in the sham first group (*p_corrected_* = 0.793, Fig. 2D). This different increase further led to overall higher RT-CoV in the taVNS first group in session 2 compared to the sham first group (*p_corrected_* = 0.013).

For the subjective task focus, we observed a marginal *stimulation*-*block* interaction (X*^2^_(3)_* = 7.693, *p* = 0.053, see Fig. S1), but post-hoc comparisons revealed no significant differences between taVNS and sham (all *p*’s > 0.38). Importantly, we did not observe a three-way interaction (X*^2^_(3)_* = 6.108, *p* = 0.106), indicating that the RT effects were not due to a modulation of task focus.

Finally, the number of *in-the-zone* trials was affected by the stimulation (X*^2^_(1)_* = 5.38, *p* = 0.02). During taVNS subjects spent an average of 189 trials per block *in the zone* (*SE* = 3.17) compared to 185 (*SE* = 3.1) during sham stimulation. No other significant main effect or interaction was observed (all *p’s* > 0.08). For the discriminatory ability parameters *d’* and *A*, we observed no stimulation specific main effects or interactions (all *p*’s > 0.22), indicating that accuracy did neither improve nor decline despite faster RTs. However, as a speed-accuracy tradeoff could be masked when performance it at ceiling level, we further analyzed commission error (CE) rate, which served as behavioral outcome in previous studies (Esterman et al., 2015, 2017). Here we found no stimulation specific main effect or interaction (all *p*’s > 0.25). More details are provided in the supplementary material.

Behavioral effects were mirrored in the electrophysiological data. The fronto-central P1-N1 amplitude during Go trials (Fig. 3A), was modulated by the *stimulation* ( *^2^_(1)_* = 4.75, *p* = 0.03) which was driven by a *stimulation-session* interaction ( *^2^_(1)_* = 4.94, *p* = 0.03; Fig. 3B): In the first session, taVNS was associated with higher amplitudes (*M* = 3.73, *SE* = 0.24) compared to sham (*M* = 3, *SE* = 0.24, *t-ratio* = -2.14, *p_corrected_* = 0.077). In the second session, this difference between taVNS and sham was absent (*p_corrected_* = 0.134), indicating again a differential effect of the stimulation depending on whether taVNS or sham was applied in the first, novel session. The parietal N1-P2 amplitude at Pz exhibited a similar *stimulation-session* interaction ( *^2^_(1)_* = 6.49, *p* = 0.01). However, here we observed no difference between taVNS and sham in the first session (*p_corrected_*= 0.121) and only a weak trend towards a difference in the second session (*t-ratio* = 2.05, *p_corrected_* = 0.094) with higher amplitudes during sham (*M* = 3.42, *SE* = 0.19) than during taVNS (*M* = 2.88, *SE* = 0.19). Additional GLMMs were fitted to test whether P1-N1 and N1-P2 amplitudes during correct Go trials can serve as predictor of RTs. Results indicate that P1-N1 amplitude but not N1-P2 amplitude significantly predicted RTs but only across both sessions and order groups (Fig. S2). When repeating the analysis separately for the taVNS-first and sham-first group, results are no longer conclusive, indicating potential loss of statistical power. More detailed information is provided in the supplementary material.

The N1-P2 amplitude for correct Nogo trials was also modulated by a significant 3-way interaction (X*^2^_(1)_* = 9.54, *p* = 0.002). Post-hoc comparisons between taVNS and sham stimulation for the different block-session combinations, however, failed to reach significance (all *p*’s > 0.5). Nevertheless, participants receiving taVNS in the first session showed a significant increase in N1-P2 amplitudes from the first (*M* = 9.87, *SE* = 1.05) to the second half of the session (*M* = 12.67, *SE* = 1.05, *t-ratio* = -4.39, *p_corrected_* < 0.001). Participants receiving sham stimulation in the first session, on the other hand, showed no difference between the first (*M* = 11.17, *SE* = 1.12) and second half of session 1 (*M* = 11.35, *SE* = 1.12, *t-ratio* = - 0.28, *p_corrected_*= 0.78, Fig. 4B). In the second session, N1-P2 amplitudes did not differ between taVNS and sham stimulation or the first and second half within a stimulation condition (all *p*’s > 0.34) Nevertheless, as for the RTs and the fronto-central P1-N1 amplitude, we see evidence of differential modulations depending on whether participants received taVNS or sham in the first session.

## Discussion

In this first electrophysiological characterization of the gradCPT, we demonstrate that ERP components resembling the canonical N2 and P3 component can be observed during both, the Go and Nogo conditions despite the continuously transitioning stimuli in the gradCPT. The Go condition was further characterized by a positive deflection preceding the N2 and P3 components. On the behavioral level, we validate previous observations (Esterman et al., 2013; Fortenbaugh et al., 2015, 2017) that reaction time and discriminatory ability during the task were modulated by the zone of attention (e.g., *in the zone* vs. *out of the zone*). More importantly, we demonstrate taVNS specific effects on RT and RT variability, which were shaped by the order in which taVNS was applied. Although taVNS improved RTs within the first session and this improvement carried over to the second session ≥ 24 hours later (Fig. 2B), it also significantly increased RT variability. These results contradict a generally improving effect of taVNS on sustained attention but highlight the importance of the context with which the stimulation is applied. Electrophysiological results mirrored this pattern, with enhanced early, sensory–attentional ERP components (P1-N1 and N1–P2; Fig. 3) and block-wise increases in Nogo-related amplitudes (Fig. 4).

### Behavioral effects

The rapid, taVNS specific decrease of RTs between block 1 and 2 in the first session suggests a more efficient stimulus processing and thus an earlier recognition of Go and Nogo stimuli. The effect is likely not just a shift in the speed-accuracy-tradeoff as overall performance did not decline in a similar pattern. This is in accordance with the role of the LC-NA system in the optimization of performance in a given task by facilitating responses to task-relevant processes (Aston-Jones & Cohen, 2005). During block 1 of the first session all participants were unfamiliar with the task, except for the two short training blocks beforehand. In order to achieve optimal performance, participants thus had to learn to differentiate the Go and Nogo stimuli as well as the appropriate stimulus-response mappings, which is crucial for optimal attentional performance (Corbetta & Shulman, 2002). This process requires more effortful top-down control provided by the dorsal attention network (Esterman et al., 2017; Rosenberg, 2026), which plays an important role in modulating activity in the visual cortex to optimize performance (Corbetta & Shulman, 2002). We assume that taVNS, applied with the onset of each stimulus, increased the phasic LC activity and thus improved the processing of task relevant information through increased attentional control from the DAN. Furthermore, this facilitation reached a relatively stable level already in the second block of the first session whereas the sham first group showed a descriptive improvement in reaction times by block 3. Thus, taVNS exerted its beneficial effects when it was applied during the initial learning phase of the task. The taVNS first group seems to have learnt the differentiation between Go and Nogo stimuli faster given that the short break between block 1 and 2 seemed to be sufficient for consolidation. The continued improvement in RTs for the taVNS first group in the second session further supports a noradrenergic modulation, as NA plays an important role in the facilitation of long-term synaptic plasticity (Sara, 2009). Both, the taVNS and the sham first group, responded faster in the second session which shows a general learning effect. However, the improvement was stronger for the taVNS first group, indicating that the stimulation amplified this naturally occurring learning effect. Participants retained the ability to differentiate faster between Go and Nogo stimuli up until the second session, which suggests lasting effects on sensory processing.

For the taVNS associated increase in RT-CoV, a different pattern emerges. In previous studies, periods of low RT variability were associated with higher activity in the Default Mode Network (DMN) and less activity in the DAN, suggesting an automatic response mode, which requires less effortful control (Esterman et al., 2013, 2014; Kucyi, Hove, et al., 2016). According to the *efficient recruitment hypothesis* (Esterman et al., 2014, 2015), top-down control areas are engaged in a more efficient and fine-tuned way during optimal sustained attention performance. Supporting this assumption, inhibiting activity through TMS in the right FEF selective increased reaction time variability and commission error rate during *in the zone* periods (Esterman et al., 2015). We observed, however, higher RT-CoV values that were systematically associated with the taVNS first group. This contradicts a generally beneficial effect of taVNS during task acquisition. While it improved learning of stimulus-response mappings during the initial part of the task, it seems to hinder an efficient, automatic response mode once task proficiency is achieved. This could be due to a continuous activation of the DAN, which was positively correlated with reaction time variability in previous studies (Esterman et al., 2014). We assume that the phasic stimulation at each stimulus onset led to a more broader activation in the DAN, which hindered an efficient automatic response mode.

Interestingly, taVNS did not affect the subjective task focus. Here we only saw a to be expected decline over time, independent of session and stimulation condition (Fig. S1). This indicates that the objective improvement in RTs was not due to participants being more focused on the task but rather due to enhanced sensory processing through allocation of attentional resources. A taVNS induced modulation of cortical arousal is also possible, which was not captured by the thought probes. In a recent study, continuous taVNS increased subjective arousal after stimulation in a sensory perception task but had no effect on salivary alpha amylase as marker of noradrenergic activity (Jelinčić et al., 2025).

Overall, the behavioral results suggest that taVNS can be beneficial when a new task requiring sustained attention is learned. Once task proficiency is achieved however, taVNS seems to be no longer helpful and to hinder automatic responding likely through continuous engagement of the DAN.

### ERPs

Regarding the ERP results, both, Go and Nogo stimuli in our study elicited a negative and a subsequent positive deflection, resembling the canonical N2 and P3 components typically observed in Go-Nogo tasks (Nieuwenhuis et al., 2003). Due to the gradual fade-in of the stimuli, these components occurred later in time compared to traditional paradigms with abrupt stimulus onsets. Across participants, the components appeared relatively stable, as indicated by the low variance in the grand average ERP. Moreover, the finding that Nogo trials elicited larger amplitudes is consistent with previous research demonstrating that N2 and P3 amplitudes tend to increase when Nogo stimuli are less frequent (Nieuwenhuis et al., 2003). The N2 component is associated with the monitoring of response conflicts between the Go and Nogo conditions (Nieuwenhuis et al., 2003) and its generator is typically located in the anterior and mid-cingulate cortex (Nieuwenhuis et al., 2003).

The P3 component, especially the more parietal located P3b, on the other side is typically associated with attention and decision making (Polich, 2007) with broader neural generators including the temporo-parietal junction (Nieuwenhuis et al., 2005). Due to its close connection to the LC-NA system (Nieuwenhuis et al., 2005; Polich, 2007), the P3 component has been investigated as potential biomarker for taVNS effects (Farmer et al., 2021; Ludwig et al., 2021). Apart from these two components, we also observed an earlier, positive deflection in correct Go trials that preceded the N2 and P3 component, similar to the Go-P2 observed by Gajewski and Falkenstein (2013). However, as most Go-Nogo studies focus on the N2 and P3 component, comparable results are sparse. In previous studies the P2 component has been linked to stimulus evaluation (Falkenstein et al., 2003; Gajewski et al., 2008; Potts, 2004) with positive correlations between P2 amplitude and reaction time variability and mean reaction time (Karamacoska et al., 2019). It has been linked to activity in the prefrontal and orbitofrontal cortex (Gajewski et al., 2008; Potts, 2004). The fact that we observed a P2 peak in the ERP for incorrect Nogo trials (commission errors) but not for correct Nogo trials (correct omissions, Fig. 3) indicates a possible missclassification into *Go* already at this early stage, as the P2 is larger for trials requiring an overt response (Gajewski et al., 2008).

Due to the nature of the gradCPT we decided to analyze peak-to-peak amplitudes, which do not depend on a baseline correction. For the Go trials we observed increased P2-N2 amplitudes only in session 1 for the taVNS-first group. In session two, these participants still exhibited numerically larger amplitudes although this difference was no longer statistically significant, mirroring in part our RT results. The association of the P2-N2 complex with stimulus evaluation and response selection (Gajewski et al., 2008) is consisted with our observation that the P2-N2 amplitude was a significant predictor of RT. The N2-P3 complex on the other hand was, as expected, larger for the Nogo condition reflecting enhanced inhibitory control and response monitoring. Here we also observed stimulation dependent effects on peak-to-peak amplitudes only in the taVNS-first group. This could be due to an increase in the P3 component as previous studies observed taVNS specific effects on P3 amplitude (Giraudier et al., 2024; Gurtubay et al., 2023). However, these ERP changes did not manifest in a behavioral effect which could indicate that this modulation was not sufficient to improve task performance in our sample. Taken together, these results indicate a noradrenergic modulation of cortical brain regions involved in attention and sensory discrimination that lasted for at least 24 hours. However, more research is needed to replicate our results and to better understand the temporal dynamics of brain activity during the gradCPT.

One limitation of our design is the lack of direct monoaminergic readouts in our study. Although much research has focused on LC-NA effects of taVNS (Ludwig et al., 2021), it likely affects other transmitter systems as well including the acetylcholine (Mridha et al., 2021), serotonine (Sclocco et al., 2019), and dopamine (Choi et al., 2024). Furthermore, we kept the stimulation intensity constant between the taVNS and sham condition to maintain comparable dosages. More recent research, however, indicates that taVNS effects on LC-NA activity depend on the subjective intensity (Vezzani et al., 2026). Stimulation at the cymba could have been perceived as more salient, thus leading to a stronger orienting response. As we did not record participants subjective ratings of stimulation intensity, this possibility cannot be ruled out. Another limiting factor is the inherently low number of Nogo trials per block, which may constrain the statistical power for detecting effects specific to response inhibition. Designs with slightly more Nogo trials (e.g., 20% Nogo instead of 10%) could help to better understand the temporal dynamics of brain activity while still eliciting a prepotent Go response. Further, as mentioned above, our sample consisted of young, healthy adults who performed with overall high accuracy. This might have limited the behavioral potential of the stimulation due to ceiling effects. More research is needed to investigate whether participants with attentional deficits could benefit more from the stimulation. Another limitation relates to the temporal stability of the observed effects. Although our design allowed us to show that effects were stable for at least 24 hours, more research is needed to understand the temporal stability over a longer period of time. In future studies, follow up session e.g., after two weeks could help to understand the long-term effects.

## Acknowledgements

This work was supported by the Data Integration Center of the University Medicine of Magdeburg. This research received no specific grant from any funding agency in the public, commercial, or not-for-profit sectors.

## Author contributionn

Conceptualization: TZ, CW; Data curation: CW; Formal analysis: CW, JPW; Funding acquisition: TZ, AG; Investigation: CW; Methodology: CW, TZ; Project administration: CW, TZ; Resources: TZ, AG; Software: CW, JPW; Supervision: TZ; Validation: CW, TZ; Visualization: CW, TZ; Writing - original draft: CW, JPW, AZ, AG, TZ;

## Data availability

EEG data and analysis code are available on OSF at osf.io/znasf/

---

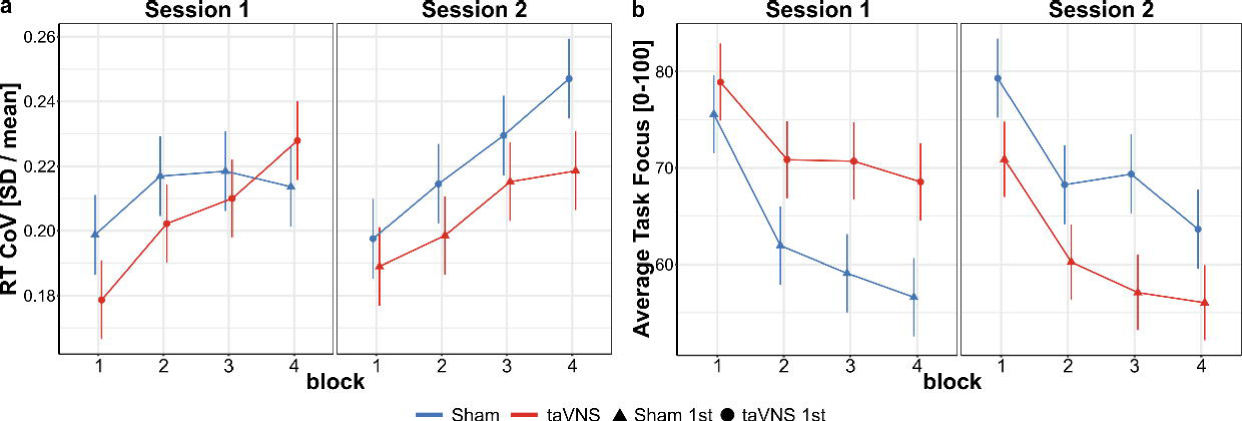

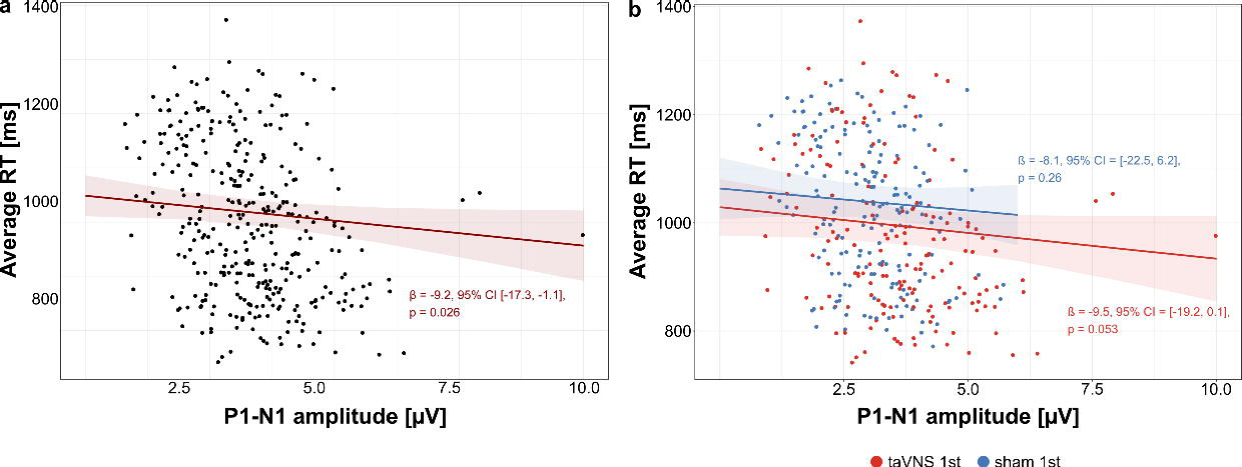

## References

1. Akkol, S., Kucyi, A., Hu, W., Zhao, B., Zhang, C., Sava-Segal, C., Liu, S., Razavi, B., Zhang, J., Zhang, K., & Parvizi, J. (2021). Intracranial Electroencephalography Reveals Selective Responses to Cognitive Stimuli in the Periventricular Heterotopias. The Journal of Neuroscience, 41(17), 3870–3878. 10.1523/JNEUROSCI.2785-20.2021

2. Aston-Jones, G., & Cohen, J. D. (2005). An integrative theory of locus coeruleus-norepinephrine function: Adaptive gain and optimal performance. Annual Review of Neuroscience, 28(1), 403–450. 10.1146/annurev.neuro.28.061604.135709

3. Barr, D. J., Levy, R., Scheepers, C., & Tily, H. J. (2013). Random effects structure for confirmatory hypothesis testing: Keep it maximal. Journal of Memory and Language, 68(3), 255–278. 10.1016/j.jml.2012.11.001

4. Bates, D., Mächler, M., Bolker, B., & Walker, S. (2015). Fitting linear mixed-effects models using Lme4. Journal of Statistical Software, 67(1), 1–48. 10.18637/jss.v067.i01

5. Bömmer, T., Schmidt, L. M., Meier, K., Kricheldorff, J., Stecher, H., Herrmann, C. S., Thiel, C. M., Janitzky, K., & Witt, K. (2024). Impact of Stimulation Duration in taVNS—Exploring Multiple Physiological and Cognitive Outcomes. Brain Sciences, 14(9), 875. 10.3390/brainsci14090875

6. Borges, U., Pfannenstiel, M., Tsukahara, J., Laborde, S., Klatt, S., & Raab, M. (2021). Transcutaneous vagus nerve stimulation via tragus or cymba conchae: Are its psychophysiological effects dependent on the stimulation area? International Journal of Psychophysiology, 161, 64–75. 10.1016/j.ijpsycho.2021.01.003

7. Borgmann, D., Rigoux, L., Kuzmanovic, B., Edwin Thanarajah, S., Münte, T. F., Fenselau, H., & Tittgemeyer, M. (2021). Technical Note: Modulation of fMRI brainstem responses by transcutaneous vagus nerve stimulation. NeuroImage, 244, 118566. 10.1016/j.neuroimage.2021.118566

8. Brainard, D. H. (1997). The Psychophysics Toolbox. Spatial Vision, 10(4), 433–436. 10.1163/156856897X00357

9. Butt, M. F., Albusoda, A., Farmer, A. D., & Aziz, Q. (2020). The anatomical basis for transcutaneous auricular vagus nerve stimulation. Journal of Anatomy, 236(4), 588–611. 10.1111/joa.13122

10. Cakmak, Y. O. (2019). Concerning Auricular Vagal Nerve Stimulation: Occult Neural Networks. Frontiers in Human Neuroscience, 13, 421. 10.3389/fnhum.2019.00421

11. Camargo, L., Pacheco-Barrios, K., Gianlorenço, A. C., Menacho, M., Choi, H., Song, J.-J., & Fregni, F. (2024). Evidence of bottom-up homeostatic modulation induced taVNS during emotional and Go/No-Go tasks. Experimental Brain Research, 242(9), 2069–2081. 10.1007/s00221-024-06876-x

12. Chamberlain, S. R., Müller, U., Blackwell, A. D., Clark, L., Robbins, T. W., & Sahakian, B. J. (2006). Neurochemical Modulation of Response Inhibition and Probabilistic Learning in Humans. Science, 311(5762), 861–863. 10.1126/science.1121218

13. Chamberlain, S. R., & Robbins, T. W. (2013). Noradrenergic modulation of cognition: Therapeutic implications. Journal of Psychopharmacology, 27(8), 694–718. 10.1177/0269881113480988

14. Chen, Y., Yang, H., Wang, F., Lu, X., & Hu, L. (2023). Modulatory effects of transcutaneous auricular vagus nerve stimulation (taVNS) on attentional processes. General Psychiatry, 36(6), e101176. 10.1136/gpsych-2023-101176

15. Choi, T.-Y., Kim, J., & Koo, J. W. (2024). Transcutaneous auricular vagus nerve stimulation in anesthetized mice induces antidepressant effects by activating dopaminergic neurons in the ventral tegmental area. Molecular Brain, 17(1), 86. 10.1186/s13041-024-01162-x

16. Cohen, M. X. (2022). A tutorial on generalized eigendecomposition for denoising, contrast enhancement, and dimension reduction in multichannel electrophysiology. NeuroImage, 247, 118809. 10.1016/j.neuroimage.2021.118809

17. Corbetta, M., & Shulman, G. L. (2002). Control of goal-directed and stimulus-driven attention in the brain. Nature Reviews Neuroscience, 3(3), 201–215. 10.1038/nrn755

18. Coull, J. T., Middleton, H. C., Robbins, T. W., & Sahakian, B. J. (1995). Clonidine and diazepam have differential effects on tests of attention and learning. Psychopharmacology, 120(3), 322–332. 10.1007/BF02311180

19. Crockett, M. J., Clark, L., Hauser, M. D., & Robbins, T. W. (2010). Serotonin selectively influences moral judgment and behavior through effects on harm aversion. Proceedings of the National Academy of Sciences, 107(40), 17433–17438. 10.1073/pnas.1009396107

20. D’Agostini, M., Burger, A. M., Jelinčić, V., Von Leupoldt, A., & Van Diest, I. (2023). Effects of transcutaneous auricular vagus nerve stimulation on P300 magnitudes and salivary alpha-amylase during an auditory oddball task. Biological Psychology, 182, 108646. 10.1016/j.biopsycho.2023.108646

21. Devoto, P., Flore, G., Saba, P., Fa, M., & Gessa, G. L. (2005). Stimulation of the locus coeruleus elicits noradrenaline and dopamine release in the medial prefrontal and parietal cortex. Journal of Neurochemistry, 92(2), 368–374. 10.1111/j.1471-4159.2004.02866.x

22. Esterman, M., Liu, G., Okabe, H., Reagan, A., Thai, M., & DeGutis, J. (2015). Frontal eye field involvement in sustaining visual attention: Evidence from transcranial magnetic stimulation. NeuroImage, 111, 542–548. 10.1016/j.neuroimage.2015.01.044

23. Esterman, M., Noonan, S. K., Rosenberg, M., & DeGutis, J. (2013). In the Zone or Zoning Out? Tracking Behavioral and Neural Fluctuations During Sustained Attention. Cerebral Cortex, 23(11), 2712–2723.

24. Esterman, M., Rosenberg, M. D., & Noonan, S. K. (2014). Intrinsic Fluctuations in Sustained Attention and Distractor Processing. The Journal of Neuroscience, 34(5), 1724–1730. 10.1523/JNEUROSCI.2658-13.2014

25. Esterman, M., & Rothlein, D. (2019). Models of sustained attention. Current Opinion in Psychology, 29, 174–180. 10.1016/j.copsyc.2019.03.005

26. Esterman, M., Thai, M., Okabe, H., DeGutis, J., Saad, E., Laganiere, S. E., & Halko, M. A. (2017). Network-targeted cerebellar transcranial magnetic stimulation improves attentional control. NeuroImage, 156, 190–198. 10.1016/j.neuroimage.2017.05.011

27. Falkenstein, M., Hoormann, J., Hohnsbein, J., & Kleinsorge, T. (2003). Short-term mobilization of processing resources is revealed in the event-related potential. Psychophysiology, 40(6), 914–923. 10.1111/1469-8986.00109

28. Farmer, A. D., Strzelczyk, A., Finisguerra, A., Gourine, A. V., Gharabaghi, A., Hasan, A., Burger, A. M., Jaramillo, A. M., Mertens, A., Majid, A., Verkuil, B., Badran, B. W., Ventura-Bort, C., Gaul, C., Beste, C., Warren, C. M., Quintana, D. S., Hämmerer, D., Freri, E., … Koenig, J. (2021). International Consensus Based Review and Recommendations for Minimum Reporting Standards in Research on Transcutaneous Vagus Nerve Stimulation (Version 2020). Frontiers in Human Neuroscience, 14, 568051. 10.3389/fnhum.2020.568051

29. Fischer, R., Ventura-Bort, C., Hamm, A., & Weymar, M. (2018). Transcutaneous vagus nerve stimulation (tVNS) enhances conflict-triggered adjustment of cognitive control. *Cognitive, Affective*, & Behavioral Neuroscience, 18(4), 680–693. 10.3758/s13415-018-0596-2

30. Folstein, J. R., & Van Petten, C. (2008). Influence of cognitive control and mismatch on the N2 component of the ERP: A review. Psychophysiology, 45(1), 152–170. 10.1111/j.1469-8986.2007.00602.x

31. Fortenbaugh, F. C., DeGutis, J., & Esterman, M. (2017). Recent theoretical, neural, and clinical advances in sustained attention research: Sustained attention. Annals of the New York Academy of Sciences, 1396(1), 70–91. 10.1111/nyas.13318

32. Fortenbaugh, F. C., DeGutis, J., Germine, L., Wilmer, J. B., Grosso, M., Russo, K., & Esterman, M. (2015). Sustained Attention Across the Life Span in a Sample of 10,000: Dissociating Ability and Strategy. Psychological Science, 26(9), 1497–1510. 10.1177/0956797615594896

33. Fortenbaugh, F. C., Rothlein, D., McGlinchey, R., DeGutis, J., & Esterman, M. (2018). Tracking behavioral and neural fluctuations during sustained attention: A robust replication and extension. NeuroImage, 171, 148–164. 10.1016/j.neuroimage.2018.01.002

34. Fox, J., & Weisberg, S. (2019). An R companion to applied regression (3rd ed.). Sage.

35. Fox, M. D., Corbetta, M., Snyder, A. Z., Vincent, J. L., & Raichle, M. E. (2006). Spontaneous neuronal activity distinguishes human dorsal and ventral attention systems. Proceedings of the National Academy of Sciences, 103(26), 10046–10051. 10.1073/pnas.0604187103

36. Gajewski, P. D., & Falkenstein, M. (2013). Effects of task complexity on ERP components in Go/Nogo tasks. International Journal of Psychophysiology, 87(3), 273–278. 10.1016/j.ijpsycho.2012.08.007

37. Gajewski, P. D., Stoerig, P., & Falkenstein, M. (2008). ERP—Correlates of response selection in a response conflict paradigm. Brain Research, 1189, 127–134. 10.1016/j.brainres.2007.10.076

38. Giraudier, M., Ventura-Bort, C., & Weymar, M. (2024). Effects of Transcutaneous Auricular Vagus Nerve Stimulation on the P300: Do Stimulation Duration and Stimulation Type Matter? Brain Sciences, 14(690). 10.3390/brainsci14070690

39. Gurtubay, I. G., Perez-Rodriguez, D. R., Fernandez, E., Librero-Lopez, J., Calvo, D., Bermejo, P., Pinin-Osorio, C., & Lopez, M. (2023). Immediate effects and duration of a short and single application of transcutaneous auricular vagus nerve stimulation on P300 event related potential. Frontiers in Neuroscience, 17, 1096865. 10.3389/fnins.2023.1096865

40. Haslacher, D., Nasr, K., Robinson, S. E., Braun, C., & Soekadar, S. R. (2021). Stimulation artifact source separation (SASS) for assessing electric brain oscillations during transcranial alternating current stimulation (tACS). NeuroImage, 228, 117571. 10.1016/j.neuroimage.2020.117571

41. Huang, R., & Clewett, D. (2024). The locus coeruleus: Where cognitive and emotional processing meet the eye. In Modern Pupillometry (pp. 3–75). Springer Nature.

42. Hulsey, D. R., Riley, J. R., Loerwald, K. W., Rennaker, R. L., Kilgard, M. P., & Hays, S. A. (2017). Parametric characterization of neural activity in the locus coeruleus in response to vagus nerve stimulation. Experimental Neurology, 289, 21–30. 10.1016/j.expneurol.2016.12.005

43. Jelinčić, V., D’Agostini, M., Ventura-Bort, C., Cascio, L., Gorianskaia, E., Weymar, M., Torta, D. M., Van Diest, I., & Von Leupoldt, A. (2025). Continuous Transcutaneous Auricular Vagus Nerve Stimulation Increases Long-Latency Neural Processing in Multiple Sensory Modalities. Psychophysiology, 62(3), e70048. 10.1111/psyp.70048

44. Karamacoska, D., Barry, R. J., De Blasio, F. M., & Steiner, G. Z. (2019). EEG-ERP dynamics in a visual Continuous Performance Test. International Journal of Psychophysiology, 146, 249–260. 10.1016/j.ijpsycho.2019.08.013

45. Keute, M., Barth, D., Liebrand, M., Heinze, H.-J., Kraemer, U., & Zaehle, T. (2020). Effects of Transcutaneous Vagus Nerve Stimulation (tVNS) on Conflict-Related Behavioral Performance and Frontal Midline Theta Activity. Journal of Cognitive Enhancement, 4(2), 121–130. 10.1007/s41465-019-00152-5

46. Kleiner, M., Brainard, D., Pelli, D., Ingling, A., Murray, R., Broussard, C., & Cornelissen, F. (2007). What’s new in Psychtoolbox-3? Perception, 36(4).

47. Kucyi, A., Esterman, M., Riley, C. S., & Valera, E. M. (2016). Spontaneous default network activity reflects behavioral variability independent of mind-wandering. Proceedings of the National Academy of Sciences, 113(48), 13899–13904. 10.1073/pnas.1611743113

48. Kucyi, A., Hove, M. J., Esterman, M., Hutchison, R. M., & Valera, E. M. (2016). Dynamic Brain Network Correlates of Spontaneous Fluctuations in Attention. *Cerebral Cortex*, bhw029. 10.1093/cercor/bhw029

49. Langner, R., & Eickhoff, S. B. (2013). Sustaining attention to simple tasks: A meta-analytic review of the neural mechanisms of vigilant attention. Psychological Bulletin, 139(4), 870–900. 10.1037/a0030694

50. Lenth, R. V. (2022). *Emmeans: Estimated marginal means, aka least-squares means* [Manual].

51. Liebe, T., Kaufmann, J., Hämmerer, D., Betts, M., & Walter, M. (2022). In vivo tractography of human locus coeruleus—relation to 7T resting state fMRI, psychological measures and single subject validity. Molecular Psychiatry. 10.1038/s41380-022-01761-x

52. Liu, K. Y., Acosta-Cabronero, J., Cardenas-Blanco, A., Loane, C., Berry, A. J., Betts, M. J., Kievit, R. A., Henson, R. N., Düzel, E., Howard, R., & Hämmerer, D. (2019). In vivo visualization of age-related differences in the locus coeruleus. Neurobiology of Aging, 74, 101–111. 10.1016/j.neurobiolaging.2018.10.014

53. Lo, S., & Andrews, S. (2015). To transform or not to transform: Using generalized linear mixed models to analyse reaction time data. Frontiers in Psychology, 6. 10.3389/fpsyg.2015.01171

54. Ludwig, M., Wienke, C., Betts, M. J., Zaehle, T., & Hämmerer, D. (2021). Current challenges in reliably targeting the noradrenergic locus coeruleus using transcutaneous auricular vagus nerve stimulation (taVNS). Autonomic Neuroscience, 236. 10.1016/j.autneu.2021.102900

55. MacDonald, S. W. S., Nyberg, L., & Bäckman, L. (2006). Intra-individual variability in behavior: Links to brain structure, neurotransmission and neuronal activity. Trends in Neurosciences, 29(8), 474–480. 10.1016/j.tins.2006.06.011

56. Manta, S., El Mansari, M., Debonnel, G., & Blier, P. (2013). Electrophysiological and neurochemical effects of long-term vagus nerve stimulation on the rat monoaminergic systems. International Journal of Neuropsychopharmacology, 16(2), 459–470. 10.1017/S1461145712000387

57. Marshall, C. A., Brodnik, Z. D., Mortensen, O. V., Reith, M. E. A., Shumsky, J. S., Waterhouse, B. D., España, R. A., & Kortagere, S. (2019). Selective activation of Dopamine D3 receptors and norepinephrine transporter blockade enhances sustained attention. Neuropharmacology, 148, 178–188. 10.1016/j.neuropharm.2019.01.003

58. McClaskey, C. M., Dias, J. W., Dubno, J. R., & Harris, K. C. (2018). Reliability of Measures of N1 Peak Amplitude of the Compound Action Potential in Younger and Older Adults. Journal of Speech, Language, and Hearing Research, 61(9), 2422–2430. 10.1044/2018_jslhr-h-18-0097

59. Mridha, Z., de Gee, J. W., Shi, Y., Alkashgari, R., Williams, J., Suminski, A., Ward, M. P., Zhang, W., & McGinley, M. J. (2021). Graded recruitment of pupil-linked neuromodulation by parametric stimulation of the vagus nerve. Nature Communications, 12(1), 1539. 10.1038/s41467-021-21730-2

60. Murphy, P. R., Robertson, I. H., Balsters, J. H., & O’connell, R. G. (2011). Pupillometry and P3 index the locus coeruleus-noradrenergic arousal function in humans: Indirect markers of locus coeruleus activity. Psychophysiology, 48(11), 1532–1543. 10.1111/j.1469-8986.2011.01226.x

61. Nieuwenhuis, S., Aston-Jones, G., & Cohen, J. D. (2005). Decision making, the P3, and the locus coeruleus–norepinephrine system. Psychological Bulletin, 131(4), 510–532. 10.1037/0033-2909.131.4.510

62. Nieuwenhuis, S., Yeung, N., Van Den Wildenberg, W., & Ridderinkhof, K. R. (2003). Electrophysiological correlates of anterior cingulate function in a go/no-go task: Effects of response conflict and trial type frequency. *Cognitive, Affective*, & Behavioral Neuroscience, 3(1), 17–26. 10.3758/CABN.3.1.17

63. Oostenveld, R., Fries, P., Maris, E., & Schoffelen, J.-M. (2011). FieldTrip: Open Source Software for Advanced Analysis of MEG, EEG, and Invasive Electrophysiological Data. Computational Intelligence and Neuroscience, 2011, 1–9. 10.1155/2011/156869

64. Pelli, D. G. (1997). The VideoToolbox software for visual psychophysics: Transforming numbers into movies. Spatial Vision, 10(4), 437–442. 10.1163/156856897X00366

65. Pervaz, I., Thurn, L., Vezzani, C., Kaluza, L., Kühnel, A., & Kroemer, N. B. (2025). Does transcutaneous auricular vagus nerve stimulation alter pupil dilation? A living Bayesian meta-analysis. Brain Stimulation, 18(2), 148–157. 10.1016/j.brs.2025.01.022

66. Pihlaja, M., Failla, L., Peräkylä, J., & Hartikainen, K. M. (2020). Reduced Frontal Nogo-N2 With Uncompromised Response Inhibition During Transcutaneous Vagus Nerve Stimulation—More Efficient Cognitive Control? Frontiers in Human Neuroscience, 14, 561780. 10.3389/fnhum.2020.561780

67. Polich, J. (2007). Updating P300: An integrative theory of P3a and P3b. Clinical Neurophysiology, 118(10), 2128–2148. 10.1016/j.clinph.2007.04.019

68. Potts, G. F. (2004). An ERP index of task relevance evaluation of visual stimuli. Brain and Cognition, 56(1), 5–13. 10.1016/j.bandc.2004.03.006

69. R Core Team. (2024). R: A language and environment for statistical computing [Manual]. R Foundation for Statistical Computing.

70. Raedt, R., Clinckers, R., Mollet, L., Vonck, K., El Tahry, R., Wyckhuys, T., De Herdt, V., Carrette, E., Wadman, W., Michotte, Y., Smolders, I., Boon, P., & Meurs, A. (2011). Increased hippocampal noradrenaline is a biomarker for efficacy of vagus nerve stimulation in a limbic seizure model: VNS, noradrenaline and seizure suppression. Journal of Neurochemistry, 117(3), 461–469. 10.1111/j.1471-4159.2011.07214.x

71. Rajkowski, J., Kubiak, P., & Aston-Jones, G. (1994). Locus coeruleus activity in monkey: Phasic and tonic changes are associated with altered vigilance. Brain Research Bulletin, 35(5-6), 607–616. 10.1016/0361-9230(94)90175-9

72. Rangon, C.-M. (2018). Reconsidering Sham in Transcutaneous Vagus Nerve Stimulation studies. Clinical Neurophysiology, 129(11), 2501–2502. 10.1016/j.clinph.2018.08.027

73. Rosenberg, M. D. (2026). A Temporal Hierarchy of Sustained Attention Dynamics. Current Directions in Psychological Science, 35(1), 3–9. 10.1177/09637214251342976

74. RStudio Team. (2021). *RStudio: Integrated development environment for R* [Manual]. RStudio, PBC.

75. Rufener, K. S., Wienke, C., Salanje, A., Haghikia, A., & Zaehle, T. (2023). Effects of transcutaneous auricular vagus nerve stimulation paired with tones on electrophysiological markers of auditory perception. Brain Stimulation, 16(4), 982–989. 10.1016/j.brs.2023.06.006

76. Ruffoli, R., Giorgi, F. S., Pizzanelli, C., Murri, L., Paparelli, A., & Fornai, F. (2011). The chemical neuroanatomy of vagus nerve stimulation. Journal of Chemical Neuroanatomy, 42(4), 288–296. 10.1016/j.jchemneu.2010.12.002

77. Sara, S. J. (2009). The locus coeruleus and noradrenergic modulation of cognition. Nature Reviews Neuroscience, 10(3), 211–223. 10.1038/nrn2573

78. Sclocco, R., Garcia, R. G., Kettner, N. W., Isenburg, K., Fisher, H. P., Hubbard, C. S., Ay, I., Polimeni, J. R., Goldstein, J., Makris, N., Toschi, N., Barbieri, R., & Napadow, V. (2019). The influence of respiration on brainstem and cardiovagal response to auricular vagus nerve stimulation: A multimodal ultrahigh-field (7T) fMRI study. Brain Stimulation, 12(4), 911–921. 10.1016/j.brs.2019.02.003

79. Smith, A., & Nutt, D. (1996). Noradrenaline and attention lapses. Nature, 380(6572), 291–291. 10.1038/380291a0

80. Szabadi, E. (2013). Functional neuroanatomy of the central noradrenergic system. Journal of Psychopharmacology, 27(8), 659–693. 10.1177/0269881113490326

81. Unsworth, N., & Robison, M. K. (2016). Pupillary correlates of lapses of sustained attention. *Cognitive, Affective*, & Behavioral Neuroscience, 16(4), 601–615. 10.3758/s13415-016-0417-4

82. Unsworth, N., & Robison, M. K. (2017). A locus coeruleus-norepinephrine account of individual differences in working memory capacity and attention control. Psychonomic Bulletin & Review, 24(4), 1282–1311. 10.3758/s13423-016-1220-5

83. Unsworth, N., & Robison, M. K. (2018). Tracking arousal state and mind wandering with pupillometry. *Cognitive, Affective*, & Behavioral Neuroscience, 18(4), 638–664. 10.3758/s13415-018-0594-4

84. Vezzani, C., Breakspear, R.-M., Thurn, L., Ettinger, U., Kühnel, A., & Kroemer, N. B. (2026). Effects of non-invasive vagus nerve stimulation on pupil dilation are dependent on sensory matching. iScience, 29(3), 114795. 10.1016/j.isci.2026.114795

85. Vossel, S., Geng, J. J., & Fink, G. R. (2014). Dorsal and Ventral Attention Systems. The Neuroscientist, 20(2), 150–159. 10.1177/1073858413494269

86. Wagenmakers, E.-J., & Brown, S. (2007). On the linear relation between the mean and the standard deviation of a response time distribution. Psychological Review, 114(3), 830–841. 10.1037/0033-295X.114.3.830

87. Wienke, C., Grueschow, M., Haghikia, A., & Zaehle, T. (2023). Phasic, Event-Related Transcutaneous Auricular Vagus Nerve Stimulation Modifies Behavioral, Pupillary, and Low-Frequency Oscillatory Power Responses. The Journal of Neuroscience, 43(36), 6306–6319. 10.1523/JNEUROSCI.0452-23.2023

88. Woller, J. P., Menrath, D., & Gharabaghi, A. (2024). EEG denoising during transcutaneous auricular vagus nerve stimulation across simulated, phantom and human data. 10.1101/2024.05.13.593884

89. Xiao, J., Hays, J., Ehinger, K. A., Oliva, A., & Torralba, A. (2010). SUN database: Large-scale scene recognition from abbey to zoo. 2010 IEEE Computer Society Conference on Computer Vision and Pattern Recognition, 3485–3492. 10.1109/CVPR.2010.5539970

90. Yamashita, A., Rothlein, D., Kucyi, A., Valera, E. M., Germine, L., Wilmer, J., DeGutis, J., & Esterman, M. (2021). Variable rather than extreme slow reaction times distinguish brain states during sustained attention. Scientific Reports, 11(1), 14883. 10.1038/s41598-021-94161-0

91. Zhang, J., & Mueller, S. T. (2005). A note on ROC analysis and non-parametric estimate of sensitivity. Psychometrika, 70(1), 203–212. 10.1007/s11336-003-1119-8

92. Zhu, S., Liu, Q., Zhang, X., Zhou, M., Zhou, X., Ding, F., Zhang, R., Becker, B., Kendrick, K. M., & Zhao, W. (2024). Transcutaneous auricular vagus nerve stimulation enhanced emotional inhibitory control via increasing intrinsic prefrontal couplings. International Journal of Clinical and Health Psychology, 24(2), 100462. 10.1016/j.ijchp.2024.100462

93. Zuberer, A., Kucyi, A., Yamashita, A., Wu, C. M., Walter, M., Valera, E. M., & Esterman, M. (2021). Integration and segregation across large-scale intrinsic brain networks as a marker of sustained attention and task-unrelated thought. NeuroImage, 229, 117610. 10.1016/j.neuroimage.2020.117610

