## Supplementary Material for "Transcutanenous auricular vagus nerve stimulation affects sustained attention in an order-dependent manner"

Christian Wienke^1,2,3^, Joshua P. Woller^4,5^, Agnieszka Zuberer^6^, Alireza Gharabaghi^4,7,8,9^, and Tino Zaehle^1,2,10,11,✉^

^1^ Institute for Medical Psychology, Otto-von-Guericke University, Magdeburg, Germany
^2^ Departement of Neurology, Section Neuropsychology, Otto-von-Guericke University, Magdeburg, Germany
^3^ Research Group “Magdeburger Arbeitsgemeinschaft für Forschung unter Raumfahrt- und Schwerelosigkeitsbedingungen” (MARS)
^4^ Institute for Neuromodulation and Neurotechnology, University Hospital Tübingen, Germany
^5^ Max-Planck-Institute for Biological Cybernetics, Tübingen, Germany
^6^ Boston University Chobanian and Avedisian School of Medicine, Department of Psychiatry, USA
^7^ Centre for Digital Health, University Tübingen, Germany
^8^ Centre for Bionic Intelligence Tübingen Stuttgart, University Hospital Tübingen, Germany
^9^ German Centre for Mental Health (DZPG), University Hospital Tübingen, Germany
^10^ Center for Behavioral Brain Sciences (CBBS), Magdeburg, Germany
^11^ German Centre for Mental Health (DZPG), partner site Halle-Jena-Magdeburg, Germany

### Computation of behavioral parameters

*Reaction times (RTs)* were calculated as in previous studies (Esterman et al., 2013; Kucyi et al., 2016) relative to the onset of each image transition. This means that a RT of 1300 ms indicates a response when the current scene was not mixed with a preceding or succeeding scene. RTs less than 1300 ms indicate responses when the current scene was still blending with the previous one, while RTs greater than 1300 ms indicate responses when the transition into the next scene had already begun. An iterative algorithm as in Esterman et al. (2013) was used to define the RTs for each trial. A response was assigned to trial *n* when the transition was between 70% for trial *n* and 40% for trial *n+1*. For faster (<70% transition for trial *n*) or slower (>40% transition for trial *n+1*) responses or multiple button presses the algorithm assigned responses as following: The response was assigned to an adjacent trial if one of them had no response. If both adjacent trials had no response, the response was assigned to the trial closest in time unless one of them was a mountain scene. In this case, subjects were given the benefit of doubt that they correctly withhold a response. In case of multiple button presses, the fastest response was used. RTs were then used to categorize each block into periods of higher and lower attention, termed “*in the zone*” and “*out of the zone*” respectively. Therefore, we computed an individual smoothed variance time course (VTC) of RTs for each block and session, analogous to Esterman et al. (2013). Missing RTs (e.g., from correct omissions during mountain scenes) were first linearly interpolated based on the two adjacent trials before z-scoring RTs for each block. Then, the absolute difference between each trial and the average from the respective block was calculated to determine the VTC. Next, the VTC was smoothed by applying a moving Gaussian kernel with a Full-Width Half Maximum of 9 trials so that each point in the smoothed VTC represents a weighted average of the surrounding trials in the original VTC (Esterman et al., 2013). The smoothed VTC was then divided into periods of higher and lower variability based on a median split. Lower variability periods (below the median) were labeled as “in the zone” while higher variability periods (above the median) were labeled as “out of the zone” (Esterman et al., 2013). We also determined the number of *in the zone* and *out of the zone* trials based on the VTC of both concatenated sessions. Therefore, the median split was not performed separately for each block but on the combined VTCs of both sessions. This allowed us to take the predictor *stimulation* into account and analyze, whether subjects spent more trials *in the zone* in one of the sessions.

*Discriminatory ability:* Trials with incorrect omissions during city scenes were counted as omission error and converted into the false alarm rate (FAR), i.e. the proportion of go-trials were subjects erroneously withhold a response. Mountain scenes where an incorrect response was given were counted as commission errors and converted into the commission error rate (CER), i.e. the proportion of Nogo-trials where subjects erroneously gave a response. The hit rate (HR) was then calculated as $1-CER$. HR and FAR were then used to calculate *d’* as measure of discrimination ability as in previous studies (Fortenbaugh et al., 2015, 2018) for each block and attentional state.

$$d'=z\left( HR \right)-z\left( FAR \right)$$

In case of no commission errors or false alarms (i.e. HR = 1 and FAR = 0 respectively), we used standard procedures by adding half an error in the respective case (Fortenbaugh et al., 2015; Treviño et al., 2021). As this procedure is rather ambiguous, we also calculated an additional parameter of discrimination called *A* (Zhang & Mueller, 2005). The advantage of this parameter is that it does not depend on the artificial insertion of errors and an implicit normal distribution. According to Zhang & Mueller (2005), *A* was computed as in the following equation:

$$A=\left\{ \begin{matrix} .75+\frac{HR-FAR}{4}-FAR\left( 1-HR \right) & \text{if }FAR\leq0.5\leq HR \\ .75+\frac{HR-FAR}{4}-\frac{FAR}{4HR} & \text{if }FAR\leq HR<0.5 \\ .75+\frac{HR-FAR}{4}-\frac{1-HR}{4\left( 1-FAR \right)} & \text{if }0.5<FAR\leq HR \end{matrix} \right.$$

### (G)LMM formulas

*Reaction times:*

$$RT\sim1+zone+stimulation*block*session+\left( 1+stimulation|subjectID \right)$$

*Reaction times coefficient of variation:*

$$RTCoV\sim1+stimulation*block*session+\left( 1+stimulation|subjectID \right)$$

*Subjective task focus:*

$$foucs\sim1+stimulation*block*session+\left( 1+stimulation+session|subid \right)$$

*Number of in the zone trials:*

$$N_{\text{ inzone }}\sim stimulation*block*session+\left( 1|subjectID \right)$$

*d’:*

$$d'\sim1+zone+stimulation*block*session+\left( 1+stimulation|subjectID \right)$$

*A:*

$$A\sim1+zone+stimulation*block*session+\left( 1+session|subjectID \right)$$

*Commission Error Rate:*

$$CE.rate\sim1+zone+stimulation*block*session+\left( 1+session|subjectID \right)$$

*Peak-to-peak amplitudes:*

$$P2P amp \sim1+stimulation*block*session+\left( 1+stimulation|subjectID \right)$$

*Peak-to-peak amplitudes for incorrect Nogo:*

$$P2P amp \sim1+stimulation*session+\left( 1|subjectID \right)$$

### Stimulation side effects

Table S1: Occurrence of aversive side effects after stimulation

| Item | Sham | taVNS | *p* |
| --- | --- | --- | --- |
| headache | 0.11 (0.39) | 0.14 (0.46) | 0.85 |
| nausea | 0.02 (0.15) | 0.02 (0.15) | 1.00 |
| dizziness | 0.07 (0.33) | 0.02 (0.15) | 0.88 |
| tingling | 0.39 (0.75) | 0.25 (0.72) | 0.35 |
| heat | 0.32 (0.64) | 0.27 (0.54) | 0.24 |
| redness | 0.59 (0.82) | 0.5 (0.66) | 0.67 |
| irritation | 0.3 (0.6) | 0.2 (0.51) | 0.43 |
| concentration | 0.45 (0.73) | 0.39 (0.58) | 0.34 |
| itching | 0.16 (0.61) | 0.11 (0.44) | 0.66 |
| Ratings were given on four-point scales between 0 (none) and 3 (strong). Values are given as average rating with standard error of the mean in parentheses. | | | |

### Causes for distraction

Table S2: Cause of distraction

| Item | Sham | taVNS | *p* |
| --- | --- | --- | --- |
| external distraction | 3.32 (1.39) | 3.52 (1.41) | 0.46 |
| task related distraction | 4.18 (1.48) | 3.77 (1.38) | 0.15 |
| mindwandering | 4.91 (1.2) | 4.77 (1.52) | 0.42 |
| Ratings were given on seven-point scales between 1 (never) to 7 (every time). Values are given as average rating with standard error of the mean in parentheses. | | | |

### Reaction Time

Table S3: Average RT on correct Go trials.

| session | block | Sham | taVNS | *p* |
| --- | --- | --- | --- | --- |
| Session 1 | 1 | 1073 (21) | 1077 (23) | 0.78 |
| Session 1 | 2 | 1073 (23) | 1020 (24) | **0.03** |
| Session 1 | 3 | 1059 (23) | 1015 (24) | **0.04** |
| Session 1 | 4 | 1051 (22) | 1008 (23) | **0.04** |
| Session 2 | 1 | 968 (23) | 1022 (26) | **0.03** |
| Session 2 | 2 | 977 (24) | 1022 (27) | **0.04** |
| Session 2 | 3 | 979 (24) | 1035 (27) | **0.03** |
| Session 2 | 4 | 986 (24) | 1024 (27) | 0.08 |
| Values represent the average RT in milliseconds estimated from the model with the standard error of the mean in parentheses. | | | | |

### Reaction time coefficient of variation

Table S4: Average RT-CoV on correct Go trials.

| session | block | Sham | taVNS | *p* |
| --- | --- | --- | --- | --- |
| Session 1 | 1 | 0.2 (0.02) | 0.21 (0.02) | 0.59 |
| Session 1 | 2 | 0.22 (0.02) | 0.24 (0.02) | 0.51 |
| Session 1 | 3 | 0.23 (0.02) | 0.25 (0.02) | 0.51 |
| Session 1 | 4 | 0.22 (0.02) | 0.28 (0.02) | **0.05** |
| Session 2 | 1 | 0.25 (0.02) | 0.19 (0.02) | 0.08 |
| Session 2 | 2 | 0.27 (0.02) | 0.2 (0.02) | **0.05** |
| Session 2 | 3 | 0.29 (0.02) | 0.22 (0.02) | **0.05** |
| Session 2 | 4 | 0.32 (0.02) | 0.23 (0.02) | **0.02** |
| Values represent the average RT-CoV estimated from the model with the standard error of the mean in parentheses. | | | | |

### Commission Error Rate

The Commission Error (CE) rate was computed as the percentage of button presses during Nogo trials separately for each block and session. Values were then analyzed using a LMM including the fixed predictors *stimulation*, *block*, and *session* up to the threeway interaction as well as the covariate *zone*. The random effects structure contained random intercepts as well as random slopes across subjects (see *(G)LMM formulas*). This analysis revealed a significant effect of *zone* ($\chi$*^2^_(1)_* = 147.35, *p* < 0.001). Similar to previous work, (Esterman et al., 2013), the CE rate was lower when participants were *in the zone* (*M:* 0.09, *SE:* 0.01) compared to being *out of the zone* (*M:* 0.19, *SE:* 0.01). No other main effects or interactions were observed (all *p*’s > 0.25). Importantly, the CE rate was similar in both, the taVNS first and sham first group in block 1 of session one (*p* = 0.51), indicating no baseline differences.

### Focus and Meta-Awareness Ratings

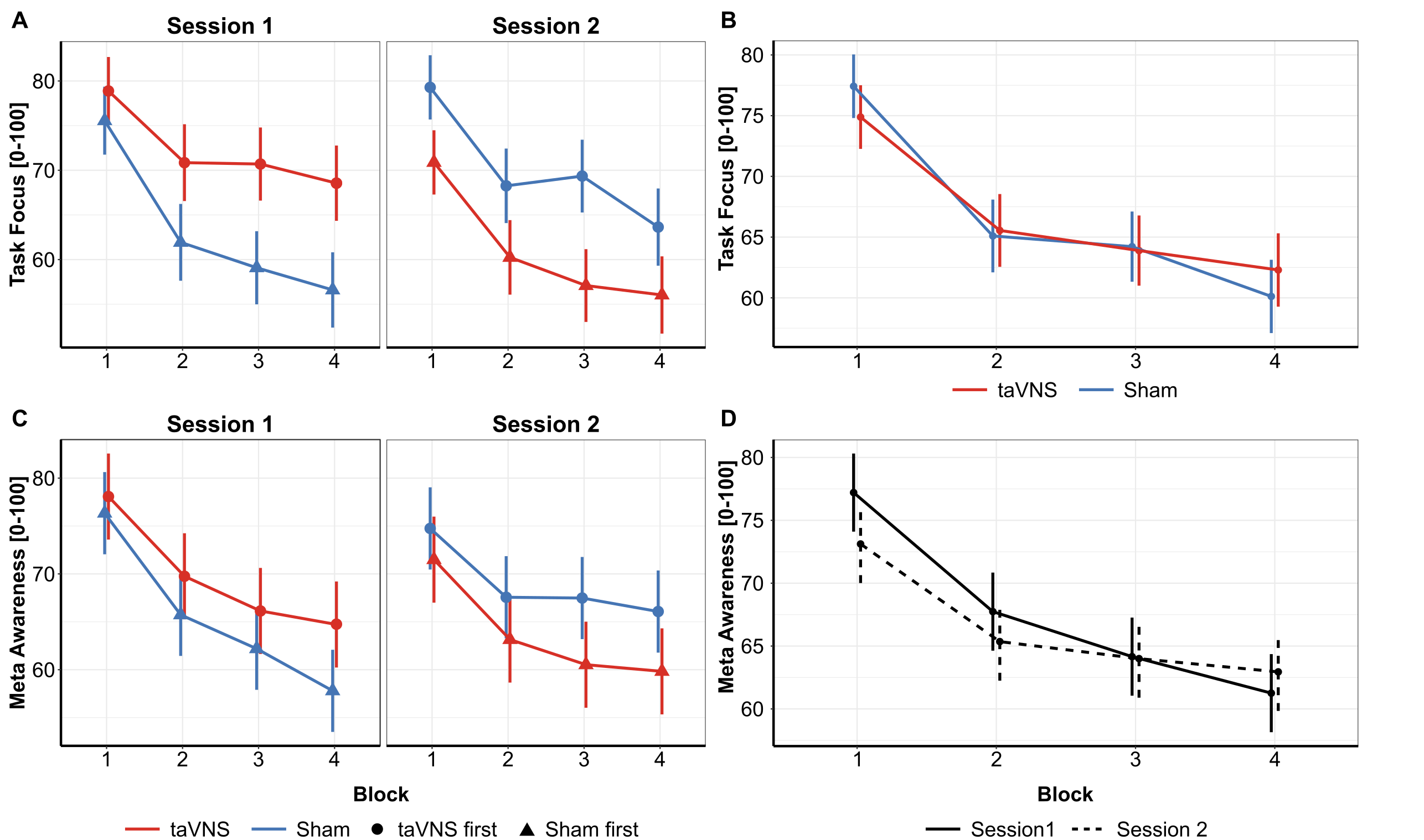

**Fig. S1. Average subjective task focus and meta awareness measured by interspersed thought probes** A) Subjective Task Focus was numerically higher in the taVNS first group, however no three-way interaction as for the RT or RT-CoV was observed. B) We only observed a marginally significant interaction between stimulation and block. Post-hoc tests, however, did not reach significance. C) For the self-reported meta awareness, a three-way interaction was again not observed. D) Here we observed a interaction between block and session, but post-hoc tests again failed to reach significance. Error bars denote s.e.m.

### Peak-to-peak amplitudes as predictor of reaction times

To connect peak-to-peak ERP amplitudes with RT, we fitted two additional GLMMs with the corresponding ERP amplitude (i.e., fronto-central P1-N1 and parietal N1-P2) as predictor of RT across sessions, blocks and stimulation conditions. This analysis revealed that the P1-N1 amplitude was a significant predictor of RT ($\chi$*^2^_(1)_* = 4.97, *p* = 0.026; Fig. S1A) while the N1-P2 amplitude was not ($\chi$*^2^_(1)_* = 0.28, *p* = 0.594; Fig. S2A). A higher fronto-central P1-N1 amplitude was associated with lower RTs ($\beta$ = -9.21, 95% CI = [-17.3, -1.11]), indicating the relevance of this component for effective stimulus processing and task performance. We then analyzed whether this connection was similarly affected by a *stimulation* - *session* interaction by fitting separate GLMMs for the taVNS-first and sham-first group. Here, we observed no significant predictive effect in the sham-first group ($\chi$*^2^_(1)_* = 1.24, *p* = 0.26) and only a marginally significant effect in the taVNS-first group ($\chi$*^2^_(1)_* = 3.74, *p* = 0.053). Again, an increased P1-N1 amplitude was associated with lower reaction times ($\beta$ = -9.53, 95% CI = [-19.2, 0.1]). However,as the 95% CI includes zero, these results need to be interpreted with care as a true effect might be absent. Further separation of the data into taVNS and sham for both, session 1 and 2 (i.e., fitting 4 GLMMs), did no longer show any predictive effect of P1-N1 amplitude on RT (all *p*’s > 0.15), indicating potential loss of statistical power.

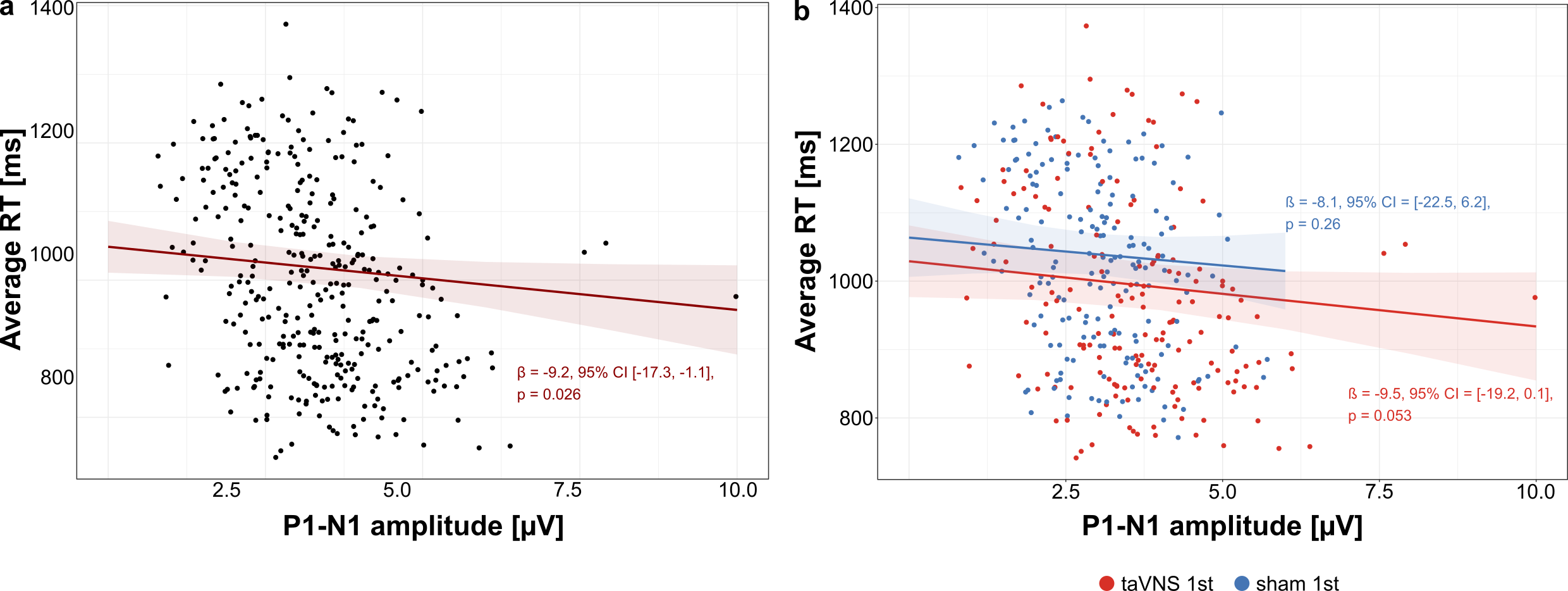

**Figure S2.** ERP amplitudes covary with reaction time. **(A)** Fronto-central (FCz + CZ) P1-N1 amplitudes as predictor of reaction time. Across all subjects, larger P1-N1 peak-to-peak amplitudes predicted faster average RTs ($\beta$ = -9.2 ms, 95% CI = [-17.3, -1.1]). **(B)** When the analysis was repeated for both participant groups, only in the taVNS-first group (red) but not in the sham-first group (blue), was the P1-N1 amplitude a marginally significant predictor of reaction times. Shaded areas represent the standard error of the regression line.

### References

Esterman, M., Noonan, S. K., Rosenberg, M., & DeGutis, J. (2013). In the Zone or Zoning Out? Tracking Behavioral and Neural Fluctuations During Sustained Attention. *Cerebral Cortex*, *23*(11), 2712–2723.

Fortenbaugh, F. C., DeGutis, J., Germine, L., Wilmer, J. B., Grosso, M., Russo, K., & Esterman, M. (2015). Sustained Attention Across the Life Span in a Sample of 10,000: Dissociating Ability and Strategy. *Psychological Science*, *26*(9), 1497–1510. <https://doi.org/10.1177/0956797615594896>

Fortenbaugh, F. C., Rothlein, D., McGlinchey, R., DeGutis, J., & Esterman, M. (2018). Tracking behavioral and neural fluctuations during sustained attention: A robust replication and extension. *NeuroImage*, *171*, 148–164. <https://doi.org/10.1016/j.neuroimage.2018.01.002>

Kucyi, A., Esterman, M., Riley, C. S., & Valera, E. M. (2016). Spontaneous default network activity reflects behavioral variability independent of mind-wandering. *Proceedings of the National Academy of Sciences*, *113*(48), 13899–13904. <https://doi.org/10.1073/pnas.1611743113>

Treviño, M., Zhu, X., Lu, Y. Y., Scheuer, L. S., Passell, E., Huang, G. C., Germine, L. T., & Horowitz, T. S. (2021). How do we measure attention? Using factor analysis to establish construct validity of neuropsychological tests. *Cognitive Research: Principles and Implications*, *6*(1), 51. <https://doi.org/10.1186/s41235-021-00313-1>

Zhang, J., & Mueller, S. T. (2005). A note on ROC analysis and non-parametric estimate of sensitivity. *Psychometrika*, *70*(1), 203–212. <https://doi.org/10.1007/s11336-003-1119-8>
